# *In vitro* Evolution of Antibody Affinity via Insertional Mutagenesis Scanning of an Entire Antibody Variable Region

**DOI:** 10.1101/2020.04.26.062786

**Authors:** Kalliopi Skamaki, Stephane Emond, Matthieu Chodorge, John Andrews, D. Gareth Rees, Daniel Cannon, Bojana Popovic, Andrew Buchanan, Ralph Minter, Florian Hollfelder

## Abstract

We report the first systematic combinatorial exploration of affinity enhancement of antibodies by insertions and deletions (InDels). Transposon-based introduction of InDels via the method TRIAD was used to generate large libraries with random in-frame InDels across the entire scFv gene that were further recombined and screened by ribosome display. Knowledge of potential insertion points from TRIAD libraries formed the basis of exploration of length and sequence diversity of novel insertions by insertional-scanning mutagenesis (ISM). An overall 256-fold affinity improvement of an anti-IL-13 antibody BAK1 as a result of InDel mutagenesis and combination with known point mutations validates this approach and suggests that the results of this InDel approach and conventional exploration of point mutations can synergize to generate antibodies with higher affinity.

**Significance:** Insertion/deletion (InDel) mutations play key roles in genome and protein evolution. Despite their prominence in evolutionary history, the potential of InDels for changing function in protein engineering by directed evolution remains unexplored. Instead point mutagenesis is widely used. Here we create antibody libraries containing InDels and demonstrate that affinity maturation can be achieved in this way, establishing an alternative to the point mutation strategies employed in all previous in vitro selections. These InDels mirror the observation of considerable length variation in loops of natural antibodies originating from the same germline genes and be combined with point mutations, making both natural sources of functional innovation available for artificial evolution in the test tube.

---

Powerful selection technologies have made *in vitro* evolution of protein binders more efficient and paved the way for the use of tailor-made antibodies in therapy. After initial selections of antibody candidates with desired specificity, lead antibodies are typically improved by affinity maturation in multiple rounds of randomisation and selection (1) to reach the sub-nanomolar affinities ideally required for targeting soluble ligands (2–4). This is usually attempted by introduction of point substitutions, either at random positions across the entire V-gene (5, 6) or in the complementary-determining regions (CDRs) e.g. by CDR walking mutagenesis (7).

In Nature, diversification of the primary antibody repertoire occurs by several mechanisms that generate variation in the regions forming the antigen-binding site, the CDRs, including considerable length variation (8–11) that is initially introduced by recombination of V(D)J gene segments. Length variations are concentrated in the CDR3 region (12), at the junctions of the joined segments, where further diversity is produced by N or P-nucleotide additions that can further extend the CDR3. The length of the CDRs considerably affects the topography of the combining site, as different shapes brought about by extension or shortening can form pockets, grooves or fill space (13, 14).

Following B-cell stimulation by the antigen, further diversification of the antigen-binding interface is generated through somatic hypermutation (SHM) (15), involving mainly point mutagenesis that preferentially targets hotspots in the CDRs (16, 17). This process is initiated by through deamination of cytosine to uracil by activation-induced cytidine deaminase (AID) leading to uracil:guanine mismatches (16). Upon removal of these uracil bases by base excision repair enzymes, error-prone DNA polymerases are then recruited to fill in the gaps and introduce mutations around the position of the deaminated cytosines. Interestingly, up to 6% of the mutations generated by SHM are insertions and deletions (InDels) (18), which occur due to misalignment of repeated DNA sequences (19, 20). Thus, insertions occur by duplication, while deletions are brought about by removal of repeated sequences (21, 22).

A small percentage of antibodies selected by *in vivo* SHM contain InDels in the CDRs 1 and 2 (1.6-6.5%) (21–24), while junctional diversity by N or P-nucleotide additions in the CDR3 confounds the analysis of SHM-derived InDels, leading to an underestimation of the total percentage of affinity-improving InDels. In other cases, introduction of InDels at random positions *in vitro* has not been possible, either because of restrictions in the diversity of InDels that could be introduced *(i.e.,* insertions by duplication in *in vitro* SHM) (22, 25) or because rational (26) or computational (27) design instead of random sampling was used. By contrast, an unusually high percentage of InDels with a functional role amongst *in vivo* affinity matured broadly neutralizing antibodies (bnAbs) to HIV-1 (28–30): ~40% of the reported anti-HIV-1 bnAbs contain InDels which accumulate during *in vivo* SHM (28). Based on the frequent occurrence of InDels among multispecific, cross-reactive antibodies one could infer that they provide a molecular solution for recognising multiple targets by providing an altered interface (enlarged or tightened), possibly even involving conformational diversity (31). The accumulation of InDels in antibodies has been attributed to extensive of *in vivo* SHM, so that even positions that are rarely modified by SHM are also altered (17, 28).

Insertions in the V-genes occur only by duplication of adjacent sequences (21, 22), so that the actual sequence diversity of the resulting insertions is limited, because they repeat existing modules. To introduce more diversity in the inserted sequences, point mutations are required in subsequent rounds of SHM. However, since the CDRs can tolerate considerable length variation, it is likely that the antibody fold can accommodate a larger number of affinity-enhancing InDels compared to those observed in antibodies affinity matured by SHM.

Affinity gains by introduction of InDels have indeed been recognised (22, 25, 26, 32, 33), but often they were by-products of campaigns focused on point mutations and not elicited systematically (32, 33). Only in *mammalian* cell surface display the action of AID leads to InDels, just as AID brings about InDels in SHM *in vivo* (22, 25). Finally, InDels have been introduced rationally based on structural analysis and natural length variation (26, 27). Taken together, only limited diversity of InDels in terms of length, position and insert sequence across the variable domains has been explored thus far.

Here we address this omission and explore libraries with in-frame indels of different lengths and high diversity of inserted sequences at *random* positions across the entire antibody variable regions (Fig. 1). We applied the new transposon-based mutagenesis approach, dubbed TRIAD (Transposition-based Random Insertion And Deletion mutagenesis) (34), to build libraries with InDels at random positions across the entire single-chain variable fragment (scFv) gene, starting with the anti-IL-13 antibody BAK1 (35), a derivative of which, tralokinumab, is under clinical investigation for asthma (36). In addition, we built libraries that explore diversity in the different *lengths* of insertions in a semi-random approach, insertional-scanning mutagenesis (ISM). These InDel libraries were starting points for antibody affinity evolution *in vitro,* leading to insertions in two loops that brought about a 256-fold affinity improvement. The observation of alternative routes to affinity maturation validate our strategy and suggest that InDel-mutagenesis can complement existing approaches.

**Fig. 1.**
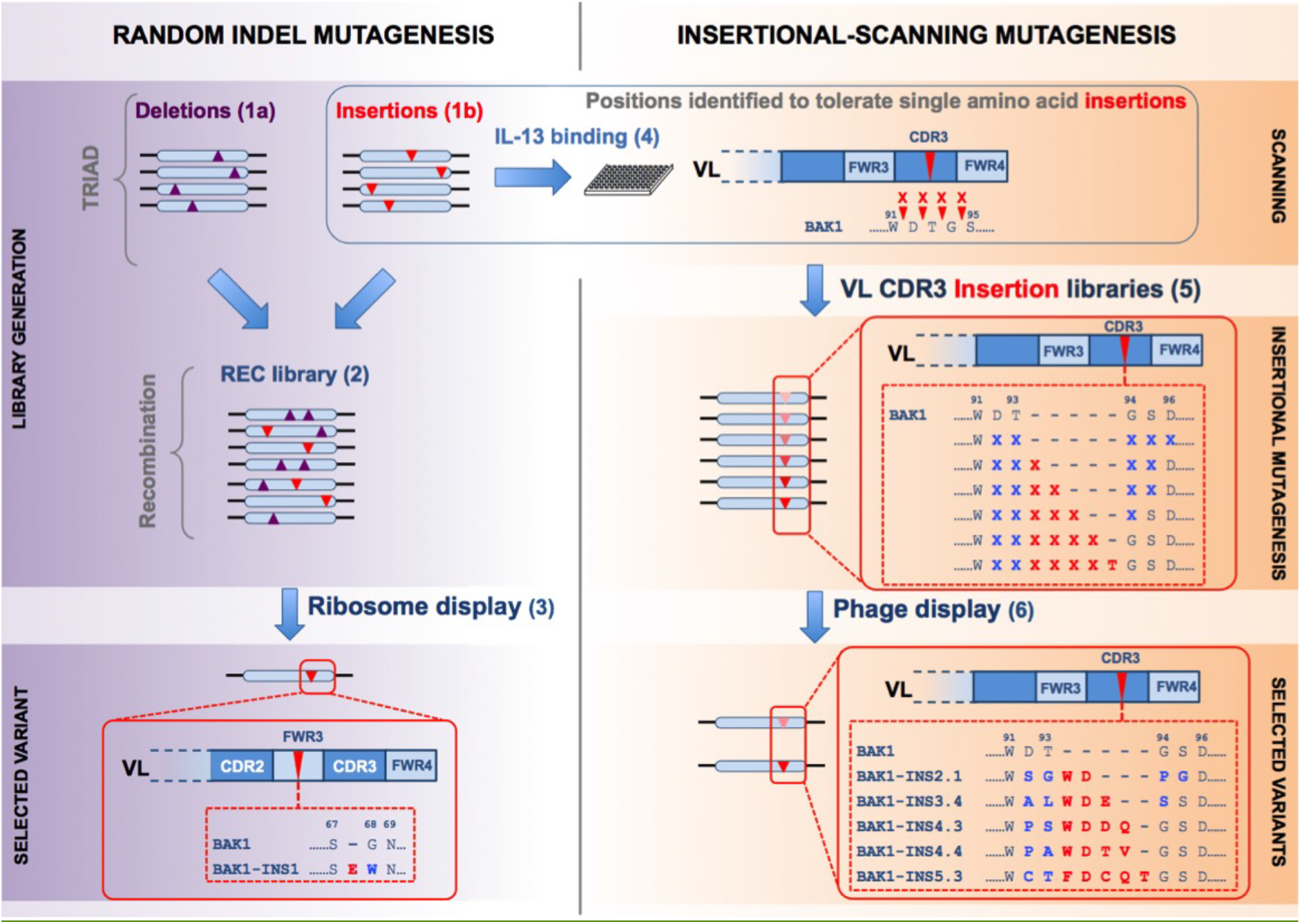
Overview of the affinity maturation of the antibody BAK1 by transposon-based TRIAD and subsequent insertional-scanning mutagenesis. TRIAD (**left column**) was applied to make libraries with deletions of 1-3 amino acids (**1a**) or single amino acid insertions (**1b**) at random positions across the scFv gene. These libraries were recombined **(2)** and four rounds of ribosome display selections for improved affinity to IL-13 were carried out by panning **(3)**. The best binder was carrying an insertion in the V_L_ FWR3 (BAK1-INS1). Scanning (**right column**) was used to guide the design of libraries with different lengths of insertions at targeted positions. A fraction of the insertion library generated in step **1b** (5,632 variants) was screened by HTRF to identify variants with insertions that retained binding to IL-13 **(4)**. Based on sequencing analysis regions able to tolerate single amino acid insertions were identified (**Fig. 4**) and the V_L_ CDR3 was chosen for targeted insertional mutagenesis. Libraries with 0-5 amino acid insertions in targeted positions in the V_L_ CDR3 were constructed **(5)**, followed by four rounds of phage display selections for improved affinity to IL-13 **(6)**.

## Results

### BAK1 affinity maturation using libraries with random InDels across the variable domains

Four libraries of the BAK1 antibody with InDels randomly distributed throughout the entire variable domains were constructed using TRIAD (34) (Figs. 1, S1 and S2): three deletion libraries (3nt-Del, 6nt-Del, 9nt-Del and 3nt-Ins, with deletions of 3, 6 and 9 nucleotides) and and one single random nucleotide triplet (NNN) insertion library (3nt-Ins) *(SI Appendix,* Table S1, Fig. 2A). Overall, the quality control of these libraries suggests that we successfully generated libraries with random in-frame InDels across the entire scFv gene, without introducing extra point substitutions, and that this custom-made diversity in sequence space can be explored in library selections.

**Fig. 2.**
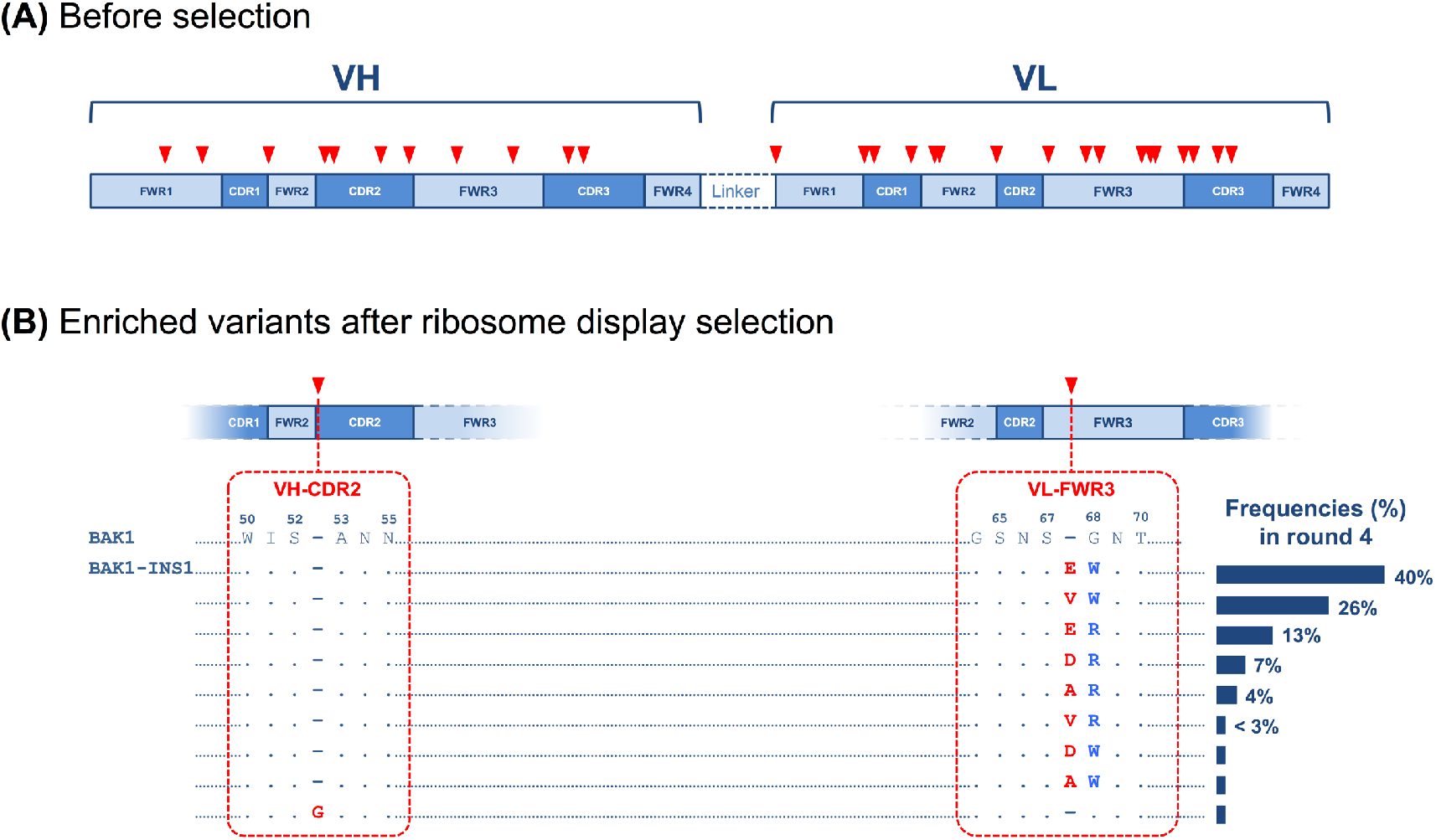
Selections of BAK1 libraries with random InDels across the scFv gene for binding to IL-13. **(A)** Prior to selection (i.e. after Step 2, Fig. 1), insertions (red) were randomly distributed across the entire V_H_ and V_L_ regions. **(B)** After selection (i.e. after Step 3 in Fig. 1) a group of variants with two consecutive mutations was enriched: an insertion (V_L_ 67a) and a point substitution (V_L_ 68), of different amino acid residues at the same position in V_L_ FWR3 (BAK1-INS variants). The predominant variant in round 4 (40%), BAK1-INS1, had an insertion (E67a) and a substitution (G68W). Sequence numbering according to Kabat (77).

To identify variants with improved affinity from these libraries, panning selections were performed by ribosome display (37), in order to take advantage of recombination before selecting from very large library sizes (>5×10^11^ variants). A shuffling library (dubbed ‘Rec’) was therefore generated by a staggered extension process (38) using the four InDel libraries of BAK1 (3nt-Del, 6nt-Del, 9-nt Del and 3nt-Ins) as templates. Sequence analysis *(SI Appendix,* Table S2) suggested that all possible BAK1 variants with single InDels and a high number of all possible BAK1 variants with two InDels per gene were present in the Rec library. Thus, the library ‘Rec’ *(SI Appendix,* Table S2) should be able to tap the potential of synergistic effects between InDels as a result of recombination of the BAK1 InDel libraries (3nt-Del, 6nt-Del, 9-nt Del and 3nt-Ins).

In order to enrich the variants with the highest affinities, four rounds of affinity-based selections of the Rec library were carried out by panning with recombinant biotinylated IL-13. Sequencing analysis of the selection outputs from rounds 1 to 4 revealed a group of variants with two consecutive mutations at the same position: a single amino acid insertion (V_L_ 67a) and accidental point substitutions (V_L_ G68X) in V_L_ FWR3 (BAK1-INS variants, Fig. 2B) were enriched over the rounds, reaching 98% of the total InDel variants in round 4 *(SI Appendix,* Fig. S4). The other in-frame indel variant that was found in round 4 (represented by the remaining 2%) contained a G52a insertion in the V_H_ CDR2, after the residue S52. The amino acid types that were enriched were insertions 67aE, 67aV, 67aD or 67aA combined with substitution G68W or G68R (Fig. 2B). One variant, BAK1-INS1, dominated the selection outcome in round 4 (40%): it has a 67aE insertion and a G68W substitution. The 67aE insertion is located within the fourth loop into the third framework region (FWR3) (Fig. 6a), sometimes referred to as CDR4 (39).

For quantitative characterisation the most enriched variant, BAK1-INS1, was converted into an IgG1 format and binding measurements revealed a 14.3-fold affinity increase (*K_D_*=140 pM) compared with the parent (*K_D_*=2,300 pM) (Table 1). No gain in affinity was observed in variants bearing either substitution G68E or G68W (BAK1-E and BAK1-W). This observation suggests that the mutation G68W was beneficial only in the presence of the 67a insertion. This observation supports the conclusion that the insertion (67a) was the seminal affinity improving feature of BAK1-INS1.

**Table 1.** Binding kinetics* and thermodynamics** of BAK1 variants (in IgG format) selected by ribosome display against IL-13.

| Loop | VH CDR 3 | VL CDR 1 | VL FWR3 | | | | $K_D$ (pM) | n-fold improvement in $K_D$ | $k_{on}$ ($M^{-1} s^{-1}$) | $k_{off}$ ($s^{-1}$) | $\Delta\Delta G$ (kcal/mol) |
| --- | --- | --- | --- | --- | --- | --- | --- | --- | --- | --- | --- |
| Residue | 99 | 27 | 67 | 67a | 68 | 69 |  |  |  |  |  |
| BAK1 | N | N | S | - | G | N | 2,300 | 1 | $5.6 \times 10^6$ | $1.3 \times 10^{-2}$ | na |
| BAK1-E | N | N | S | - | E | N | 17,100 | 0.13 | $3 \times 10^6$ | $4.4 \times 10^{-2}$ | nd |
| BAK1-W | N | N | S | - | W | N | 1,400 | 1.6 | $6 \times 10^6$ | $8.7 \times 10^{-3}$ | nd |
| BAK1.1 | S | I | S | - | G | N | 50 | 46 | $6.2 \times 10^6$ | $3.8 \times 10^{-4}$ | -3 |
| BAK1-INS1 | N | N | S | E | W | N | 140 | 14.3 | $7.1 \times 10^6$ | $1 \times 10^{-3}$ | -1 |
| BAK1-INS1.1 | S | I | S | E | W | N | 9 | 256 | $2.9 \times 10^6$ | $2.5 \times 10^{-5}$ | -4.9 |

\* $K_D$ values of the human IgG1 variants were determined by surface plasmon resonance. The n-fold improvement in $K_D$ is the ratio $K_D^{Variant} / K_D^{Parent}$ . \*\* $\Delta\Delta G$ was calculated from the equation $\Delta\Delta G = -RT \ln (K_D^{Parent} / K_D^{Variant})$ .

Although the entire BAK1 scFv had previously been scanned for affinity-improving point mutations (35), no change in affinity could be attributed to point substitutions in the V_L_ FWR3 in the present study. Furthermore, no affinity-enhancing substitutions in the V_L_ FWR3 were found (as in (35)), despite enrichment of variants with other point mutations, namely V_L_ CDR1 N27Y (BAK1.4) and V_H_ CDR3 N99S (BAK1.55) *(SI Appendix,* Table S3). These are substitutions that make up the variant BAK1.1 (identified previously by ribosome display selections of BAK1 antibody libraries (35)), suggesting that our experiments tap similar effects as a past affinity maturation campaign (and thus do not require the presence of an insertion). In our study, large affinity effects seem enabled by targeting V_L_ FWR3 with an insertion (at position 67) and not through point substitutions (Tables 1 & *SI Appendix* S3). This is the first example of a deliberate library strategy in which insertions are targeted as the main cause of antibody affinity maturation.

The sequencing analysis of variants that were enriched during round 4 (Fig. 2B), suggests that the insertion 67a was beneficial only in the presence of the specific amino acid substitution G68W. While the presence of the variant 67aE/G68 in the original insertion library was confirmed, only the 67aE/G68W variant (41%) was enriched, implying a positive synergistic effect between the two mutations. This idea is supported by the observation that similar variants combining G68W with a different residue inserted at position 67 were also enriched: 67aV/G68W and 67aA/G68W with 26% and <3% in round 4, respectively. Taken together, these results underline the importance of sampling insertions with high amino acid diversity and point substitutions in positions flanking the insertion to identify insertions with beneficial effect on affinity.

### Synergistic interaction between beneficial insertion and point substitutions

To study epistatic interactions of insertions and point substitutions on antibody affinity, we determined the combined affinity effect of the beneficial insertion in FWR3 (BAK1-INS1) and two beneficial substitutions (found in BAK1.1) (Fig. 3 and Table 1). Combination of both the insertion and substitutions resulted in an IgG variant (BAK1-INS1.1; *K_D_* = 9 pM) with higher affinity than either point substitutions (BAK1.1; *K_D_* = 50 pM) or the insertion alone (BAK1-INS1; *K_D_* = 142 pM). BAK1-INS1.1 possessed an overall 256-fold improvement in affinity compared to the parent, which is 6-fold higher than that of the substitution variant BAK1.1, with a 6.5-fold lower dissociation constant *k_off_* (2.45 × 10^-5^ s^-1^), consistent with an off-rate selection. The effect of combining insertion and point substitutions was synergistic, as the free energy change that resulted from combining insertion and point substitutions (BAK1-INS1.1; −4.9 kcal/mol) was higher than the sum (−4 kcal/mol) of the free energies of the variants with insertion (BAK1-INS1; −1 kcal/mol) and substitutions (BAK1.1; −3 kcal/mol), while for additive mutations the free energy change would be equal to the sum. These results demonstrate that beneficial insertions and point substitutions can have additive effects on affinity. This suggest a protein engineering strategy in which selections of libraries with insertions and substitutions could be performed separately and then combined to exploit synergistic effects on affinity.

**Fig. 3.**
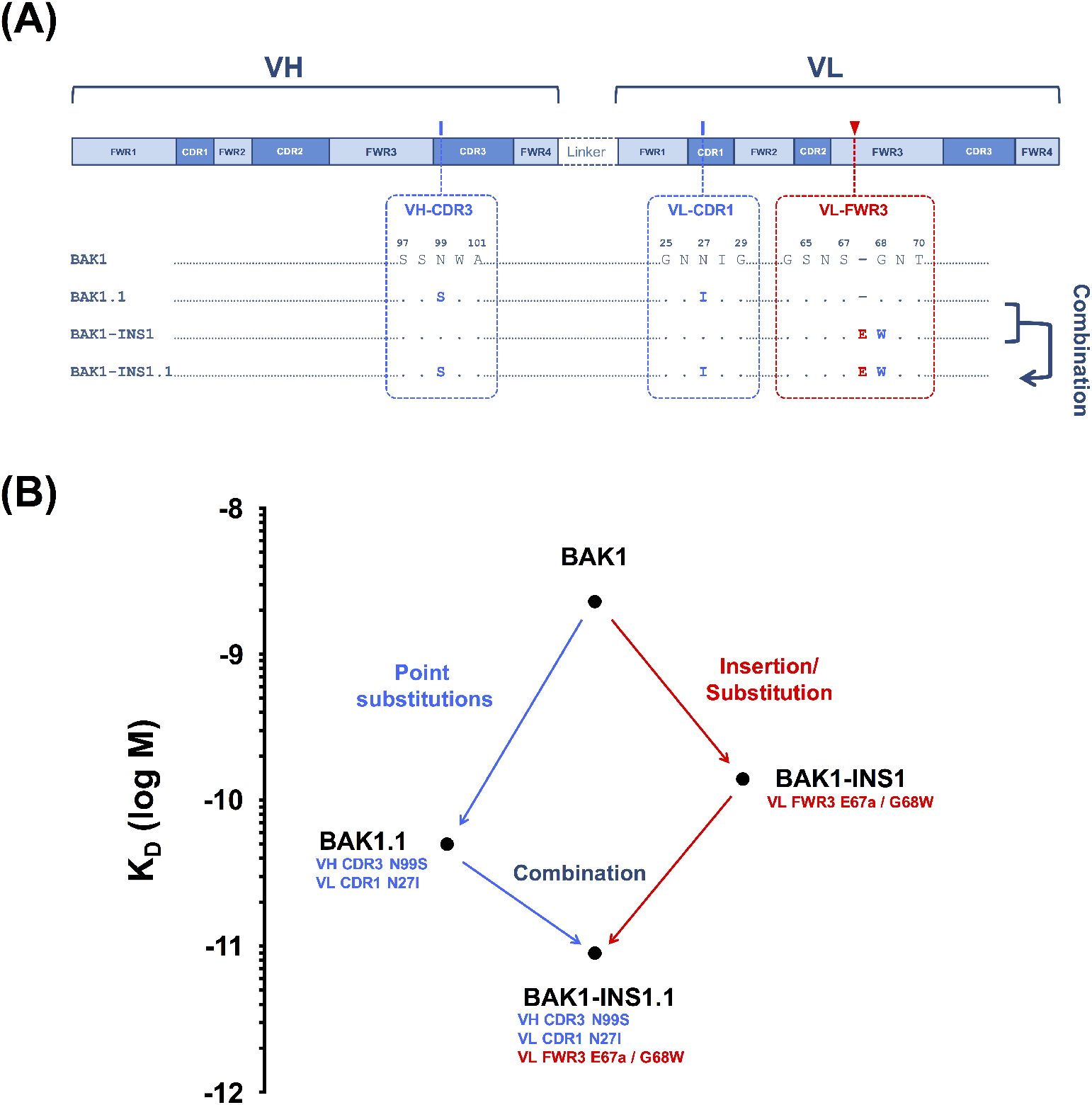
Synergy between affinity-improving substitutions in the CDRs and an insertion in V_L_-FWR3. **(A)** Mutations in the sequence of BAK1 variants: BAK1.1, a previously generated (35) variant with two beneficial substitutions (V_H_ CDR3 N99S / V_L_ CDR1 N27I); BAK1-INS1, combining two consecutive mutations, an insertion and a substitution in VL FWR3 (E67a/G68W; this study); BAK1-INS1.1, combining four mutations from both BAK1.1 and BAK1-INS1. **(B)** Mutational trajectories and improvements in affinity between parental BAK1 and variants BAK1.1, BAK1-INS1 and BAK1-INS1.1.

### BAK1 affinity maturation by insertional-scanning mutagenesis (ISM)

To identify further affinity enhancing insertions in BAK1, insertions with diverse lengths and amino acid compositions were sampled in a novel strategy. This semi-random approach, dubbed insertional scanning-mutagenesis (ISM) consisted of two stages: identification of sites in which insertions were tolerated, followed by exploration of diverse insertions in these sites. Practically, 5,632 variants of the 3nt-Ins TRIAD library (in scFv format) with single amino acid insertions (Fig. 2A, *SI Appendix* Table S1) were screened for variants with insertions that retained binding to IL-13 in a microplate-based HTRF (Homogeneous Time Resolved Fluorescence) assay. Despite the relatively low throughput, multiple insertion mutants for each position exist, so that the question of insertional tolerance could be addressed by the library outcome. Positions tolerant to insertions across the variable regions were revealed by sequencing analysis of the hits (Fig. 4), by determining the number of hits with insertion at a given position, even though different amino acid residues were inserted. Fig. 4 displays the sequence analysis of the IL-13 binders obtained in this experiment. After disregarding insertions in the N-terminus of the V_H_ (a possibly unfolded or structurally plastic terminus likely to be insertion-tolerant), 11 positions tolerating insertions in the loops V_H_ FWR3, V_H_ CDR2, V_L_ CDR3 and V_L_ FWR3 were found. Indeed, the position G68 in the V_L_ FWR3 loop (5 hits) had previously been shown to tolerate an insertion in BAK1-INS1. A cluster of four positions with high frequency was found in the V_L_ CDR3 before the residues D92, T93, G94 and S95 with 4, 10, 18 and 1 hit, respectively *(SI Appendix* Table S5). Only one position with high frequency (before A53) was observed in the V_H_ CDR2.

**Fig. 4.**
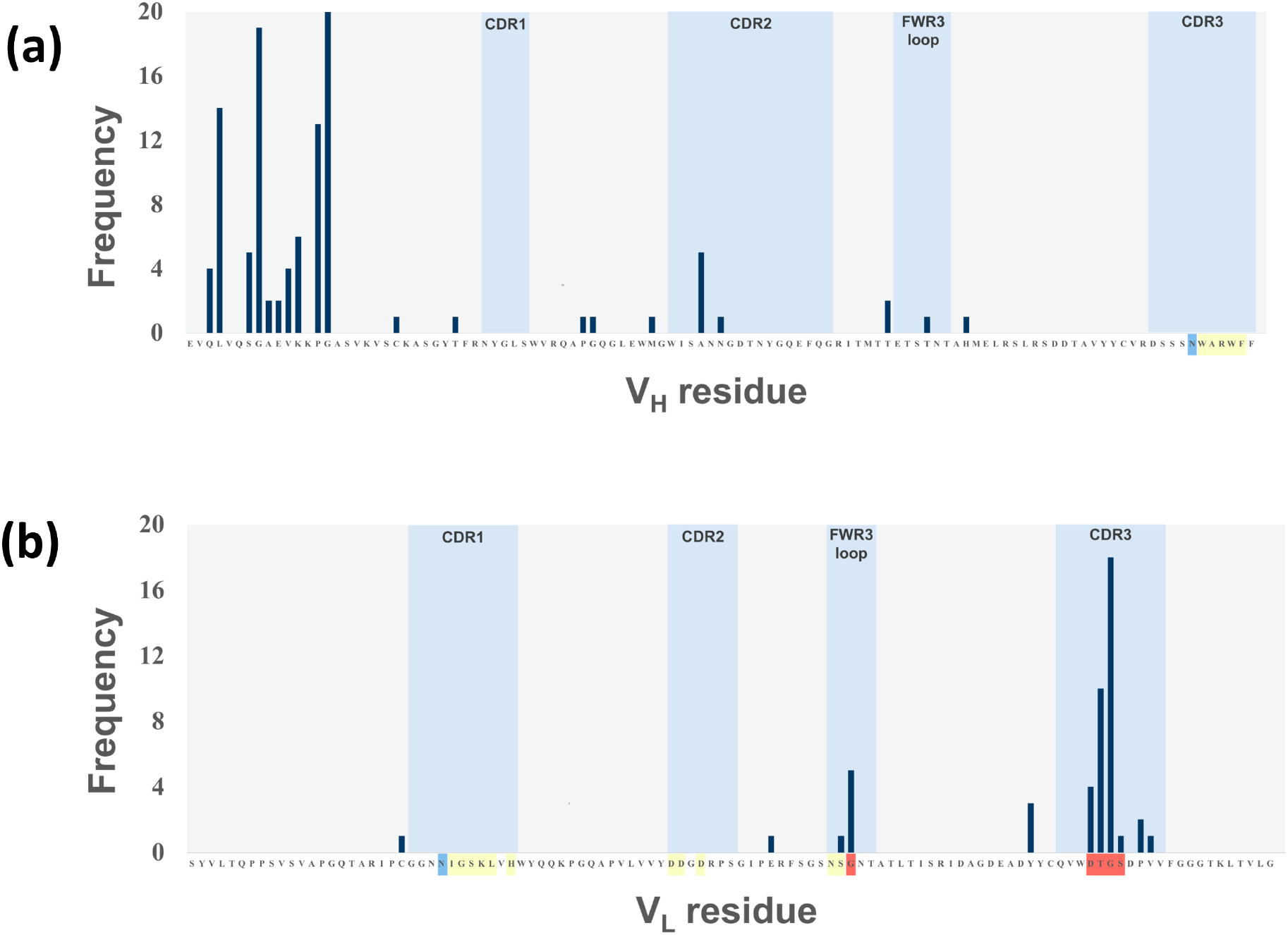
Tolerance for insertions across the variable regions of the BAK1 antibody. 5,632 variants from the TRIAD library with single amino acid insertions (Fig. 2A; *SI Appendix,* Table S1) were randomly picked and screened for retention of IL-13 binding. The positions where insertions resulted in binding levels similar or better than BAK1 are indicated by the frequency observed for insertions before the residue of the (a) V_H_ and (b) V_L_ sequence of the BAK1 antibody, respectively. The frequency refers to the number of hits at a given position (even if the inserted amino acid differs). The loop showing the highest frequency of insertion tolerance *(i.e.,* VL CDR3) was chosen as the target for insertional mutagenesis. The positions are colour-matched to Fig. 6, where the *yellow* colour indicates residues that constitute the paratope, positions shown in *blue* are those for which substitutions had been shown previously to improve affinity (in BAK1.1) (35) and positions in *red* correspond to the insertions identified in this work.

The positions tolerating insertions in the CDRs and FWR3 loop were considered possible sites to target by insertional mutagenesis, for which libraries with different lengths of insertions and with high diversity of amino acid replacements were designed and synthesised. Leaving aside 97 N-terminal insertions, 49 of the remaining 62 hits (=79%) are located in the CDRs or FWR3 loop. Amongst these 49 33 are located in V_L_ CDR3 (in positions D92 – S95 of the V_L_ CDR3; 67%), which was thus chosen as the target for saturation mutagenesis. Five libraries with 0-5 insertions as NNS codons between residues 93 and 94 and randomised flanking residues (92, 93 and 94–96) were constructed (Table 2). In total residues 5 or 6 NNS positions lead to library sizes between 10^9^-10^10^ variants

**Table 2.**
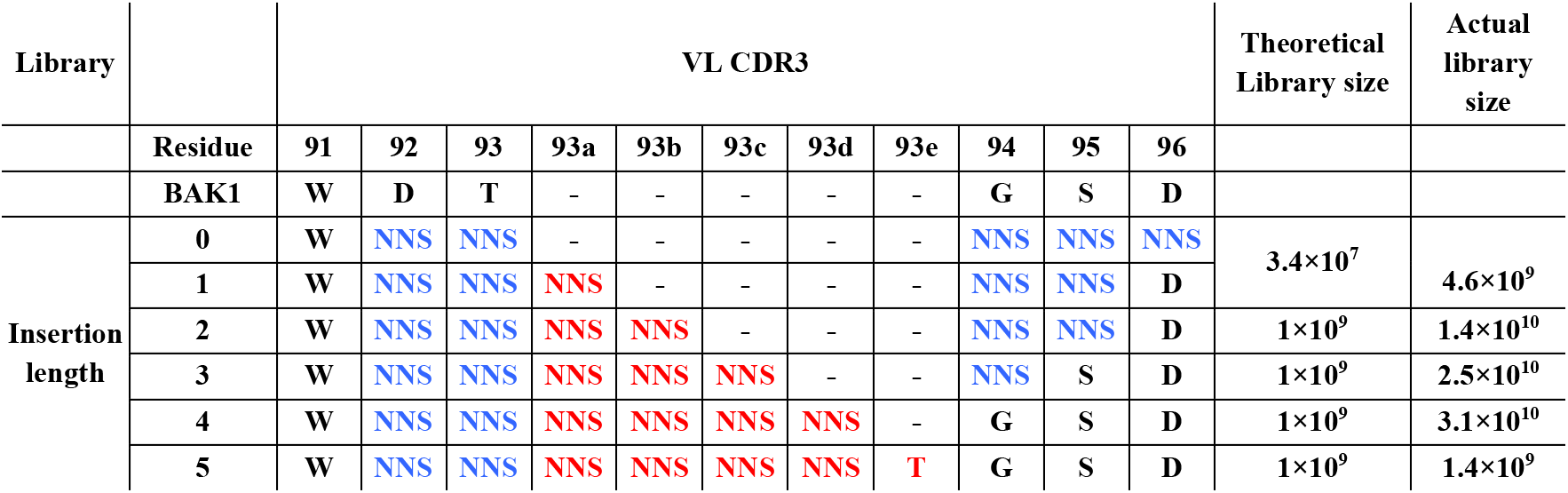
Design of libraries with different lengths of insertions in the V_L_ CDR3.

To enrich for the highest affinity variants, four rounds of KD-based phage display selections of the libraries with insertions in the V_L_ CDR3 were performed. For each library, 384 variants of the round 4 selection output *(SI Appendix,* Fig. S5) were re-screened in a competition HTRF screen for binding to human IL-13 followed by sequencing of all the hits and affinity determination in IgG format (Table 3). Analysis of the *K_D_* values of IgG variants (Table 3) showed that large improvements (> 5-fold), INS2.1, INS3.4, INS4.3, INS4.4 and INS5.3 (14-, 7.7, 26-, 68- and 48-fold improvements in *K_D_*, respectively), were associated with insertions longer than two amino acids (2 to 5 amino acids) and a motif (93aW/93bD or 93aF/93bD for the insertions of 2-4 or 5 amino acids, respectively). These data (displayed in Fig. 5a) suggest that longer inserts make it more likely to find large effects on affinity, but length alone is not a sufficient criterion for high affinity. The lengths of the V_L_ CDR3 for the parent BAK1 is 11, which is extended in insertion mutants (to 13 in INS2.1, and up to 15 in INS5.3). By contrast the V_L_ CDR3 loops of the λ-light chains observed in Nature range between 7 and 13 amino-acids (10). The successful selection of affinity-enhancing insertions with high diversity amino acid replacements stands in contrast to the selection from libraries with single or no insertion. Screening of these showed only one variant with a small improvement in affinity (INS1.6; 2.9-fold), indicating that variants without insertion in V_L_ CDR3 were hard to improve, unless an insertion in V_L_ CDR3, is present.

**Fig. 5.**
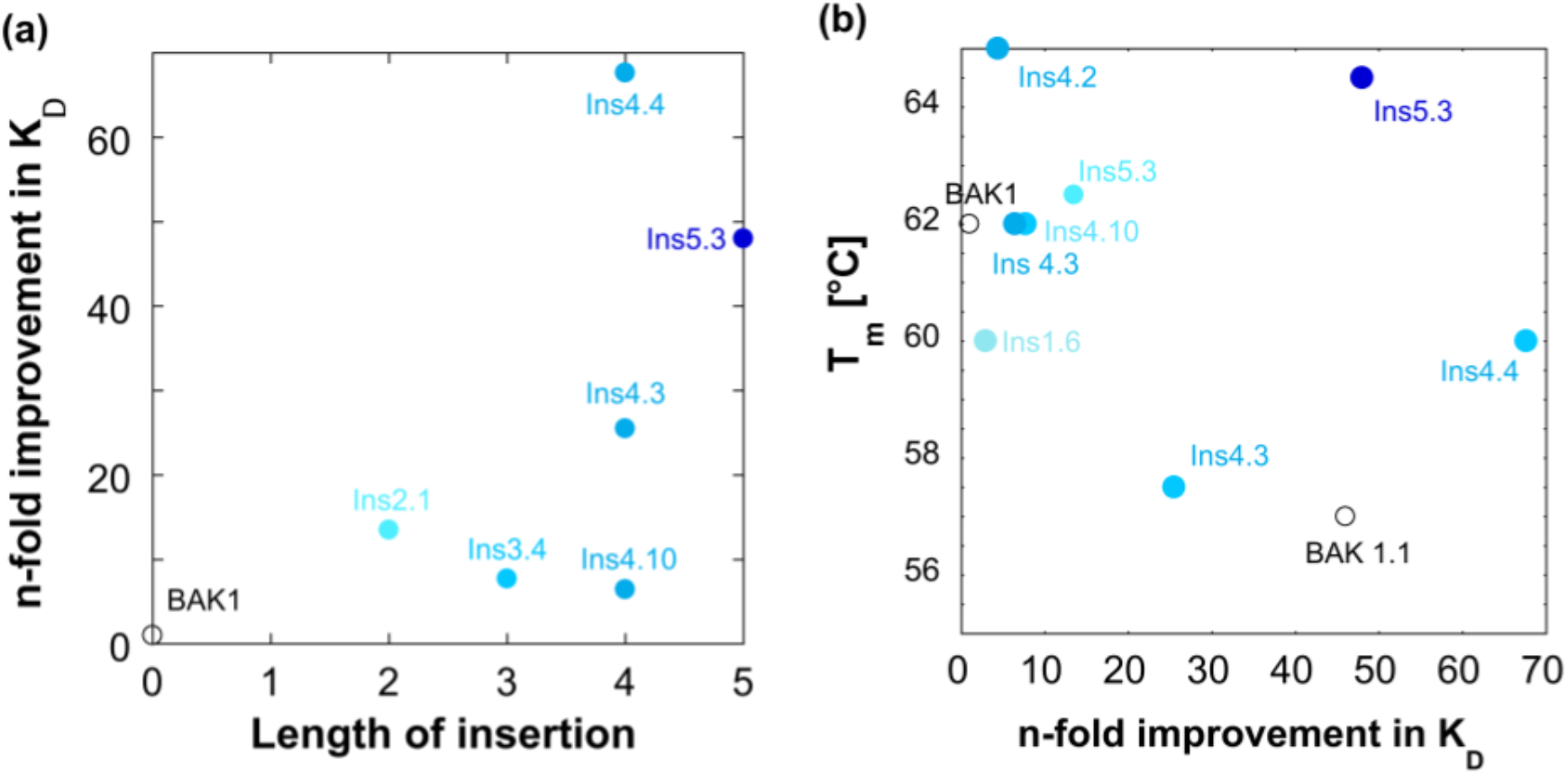
Correlations of functional and structural features with the binding enhancements. **(a)** The length of the insertion plays a partial role in affinity maturation: increasing insert sizes apparently provide more opportunity for interactions and antigen complementarity. While longer inserts generally tend to bind more tightly, the correlation also shows considerable scatter, suggesting that idiosyncrasies of specific interactions may or may not contribute. **(b)** The correlation between melting temperatures and fold improvement in *K_D_* shows a trade-off between affinity and stability for the variants with the largest improvements in affinity. There is one exception: INS 5.3 is a variant with a five-amino acid insertion, which possessed higher thermodynamic stability than the parent BAK1 (T_m_=64.5) *and* a large gain in affinity. Thermal stability measurements of improved BAK1 IgG variants with insertions in the V_L_ CDR3 were carried out by differential scanning fluorimetry (49).

**Table 3.** Binding affinity* of improved BAK1 variants with insertions in the V_L_ CDR3.

| Loop |  | VH CDR3 | VL CDR1 | VL CDR3 |  |  |  |  |  |  |  |  |  |  | Affinity |  |  |  | Thermal stability |
| --- | --- | --- | --- | --- | --- | --- | --- | --- | --- | --- | --- | --- | --- | --- | --- | --- | --- | --- | --- |
| Residue |  | 99 | 27 | 91 | 92 | 93 | 93a | 93b | 93c | 93d | 93e | 94 | 95 | 96 | K<sub>D</sub> (pM) | Fold improvement in K<sub>D</sub> | k<sub>on</sub> (10<sup>6</sup> M<sup>-1</sup> s<sup>-1</sup>) | k<sub>off</sub> (10<sup>-4</sup> s<sup>-1</sup>) | T<sub>m</sub> (°C) |
| parent | BAK1 | N | N | W | D | T | - | - | - | - | - | G | S | D | 2,300 | 1 | 4.9 | 100 | 62 |
| 0 | BAK1.1 | S | I | W | D | T | - | - | - | - | - | G | S | D | 50 | 46 | 8.5 | 4.3 | 57 |
| 1 | Ins1.6 | N | N | W | S | W | H | - | - | - | - | G | A | D | 800 | 2.9 | 6.5 | 5 | 60 |
| 2 | Ins2.1 | N | N | W | S | G | W | D | - | - | - | P | G | D | 170 | 14 | 8 | 10 | 62.5 |
| 3 | Ins3.4 | N | N | W | A | L | W | D | E | - | - | S | S | D | 300 | 7.7 | 5.7 | 10 | 62 |
| 4 | Ins4.2 | N | N | W | V | P | P | H | I | R | - | G | S | D | 530 | 4.4 | 3.1 | 10 | 65 |
|  | Ins4.3 | N | N | W | P | S | W | D | D | Q | - | G | S | D | 90 | 25.5 | 6.9 | 6.3 | 57.5 |
|  | Ins4.4 | N | N | W | P | A | W | D | T | V | - | G | S | D | 34 | 68 | 5.3 | 1.8 | 60 |
|  | Ins4.10 | N | N | W | T | T | F | D | Y | P | - | G | S | D | 360 | 6.4 | 4.1 | 10 | 62 |
| 5 | Ins5.3 | N | N | W | C | T | F | D | C | Q | T | G | S | D | 48 | 48 | 2.5 | 1.2 | 64.5 |

\* K<sub>D</sub> values of the human IgG1 variants were determined by surface plasmon resonance (T200, Biacore). The fold improvement in K<sub>D</sub> is the ratio K<sub>D</sub><sup>Variant</sup> / K<sub>D</sub><sup>Parent</sup>. The errors of the k<sub>on</sub> and k<sub>off</sub> measurements were between 0.01 and 0.09 × 10<sup>6</sup> M<sup>-1</sup>s<sup>-1</sup> and between 0.001- and 0.34 × 10<sup>-4</sup> s<sup>-1</sup>, respectively.

### Selected variants with insertions are not thermally destabilised

The perception that the stability cost of the mutational load that accompanies improvements in protein function has led to the model of trade-offs between affinity and stability in directed evolution (40, 41) that is supported by many studies (42–44). Indeed, the ‘one amino acid at a time’ model (45, 46) proposes to proceed through relatively conservative mutation regimes, to allow cycles of structural destabilisation and repair (47) and satisfy the stability requirement for clones with novel functions. It has been postulated (48) that, despite being potential sources of functional innovation, InDels are more disruptive than point mutations, implying their mutational damage to structure is higher and InDel strategies intrinsically less likely to succeed. To address this issue experimentally, we measured the thermal stability of the improved IgG variants by differential scanning fluorimetry (DSF) (49) (Fig. 5b and *SI Appendix* S6, Table 3). Most variants with small improvements in affinity (INS2.1, INS3.4, INS4.2) were as stable as the parent IgG (BAK1; T_m_=62 °C), thus the insertions did not have a negative impact on antibody stability. Most variants with large improvements in affinity had a slight loss in stability (INS4.3; T_m_=58 °C, INS4.4; T_m_=60 °C), and the same loss was observed for the variant with two point substitutions (BAK1.1; T_m_=57 °C), indicating that the trade-off between affinity and stability was observed for *both* the insertion and the substitution variants with large improvements in affinity. The only exception was the variant with a five-amino acid insertion (INS5.3), which besides conferring large gain in affinity, also possessed higher stability than the parent (T_m_ =65 °C). This improvement in stability could be attributed to the insertion of two cysteine residues that likely form a disulphide bond that could stabilize the loop. Intra-loop disulphide bonds in the V_L_ CDR3 have not been observed in natural antibodies. Thus, different effects were observed for insertions with different lengths and amino acid composition in the same region in the VL CDR3, suggesting that specific amino acid substitutions have idiosyncratic effects on function. By and large, however, in most cases a longer insert has an increased potential to exert a substantial effect on affinity.

Overall, these results show that despite the low tolerance for insertions, similar effects as point substitutions on antibody stability are observed: they can be neutral, deleterious or beneficial, being subject to affinity/stability trade-off but, according to our present data, InDels are no worse than point mutants, as recently observed in InDel mutagenesis of an enzyme (38).

## Discussion

### Insertion hotspots in the CDRs and FWR3 loops emerge from InDel evolution

This work identified insertions that brought about large affinity improvements in two different loops of the same antibody, the V_L_ CDR3 and the V_L_ FWR3 loops: (i) two to five amino acid residue insertions in the VL CDR3, replacing the residues D92 - S95 and (ii) a single amino acid insertion in the VL FWR3 loop, replacing the residue G68. Two point substitutions that improved BAK1 affinity (BAK1.1 variant) were found in different loops, the V_H_ CDR3 (N99S) and V_L_ CDR1 (N27I).

The crystal structure of tralokinumab (derived from the variant BAK1.1) in complex with human IL-13 (50) provides a basis for rationalisation of the roles of affinity-enhancing mutations with respect to the antigen binding site. Fig. 6B highlights the structural paratope evident from this structure (50), which reveals that the positions of the insertions are on the periphery of the antigen-binding site (Fig. 6), with residues G68 in V_L_ FWR3 loop and T93 in V_L_ CDR3 constituting part of the paratope.

**Fig. 6.**
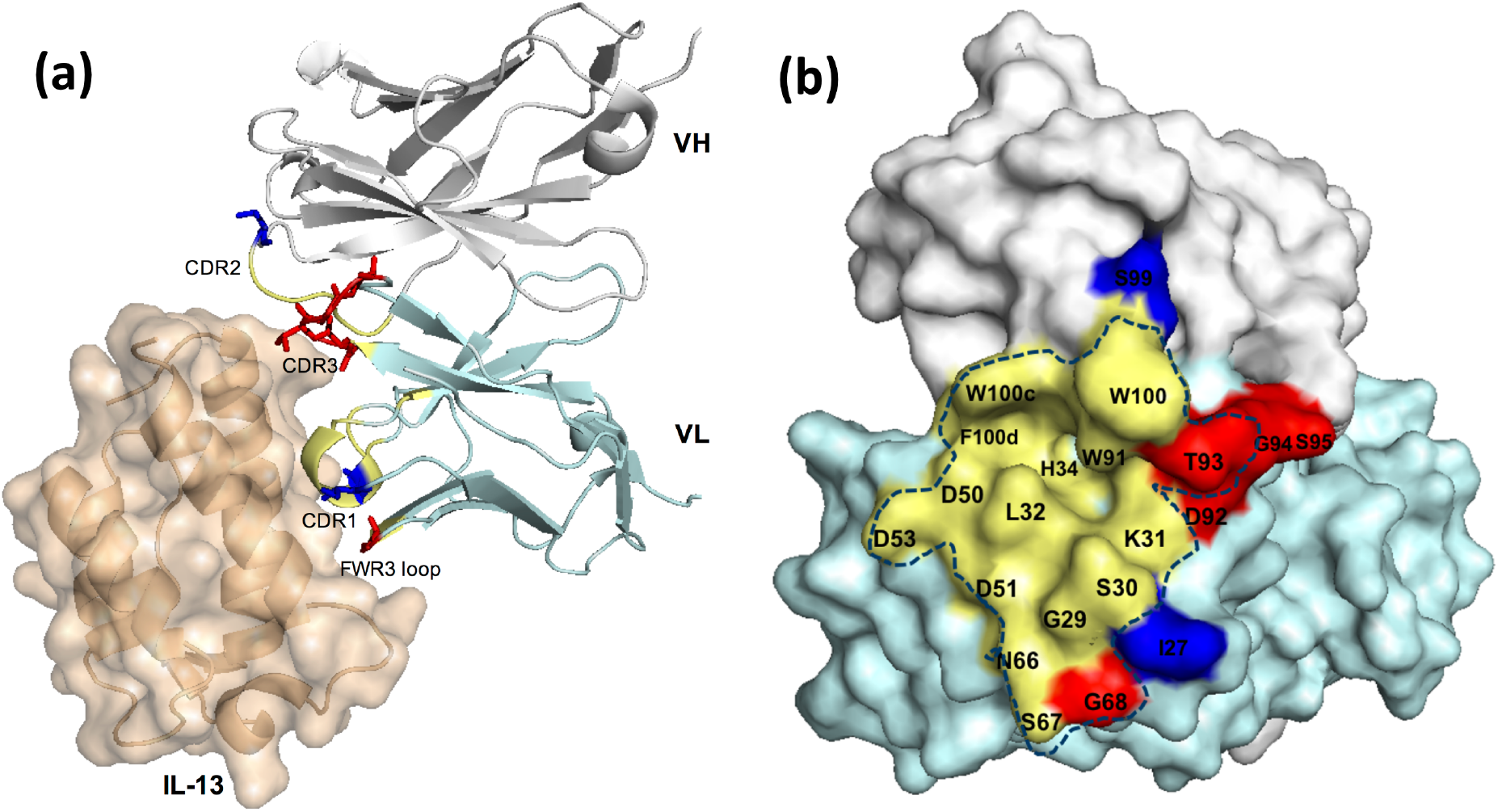
Sites of affinity-enhancing mutations modelled on the tralokinumab::IL-13 complex crystal structure (50). **(a)** Insertions with large beneficial effect on BAK1 affinity were found in two different loops, the V_L_ FWR3 loop (G68) and the V_L_ CDR3 (D92-S95), in close proximity to the antigen, IL-13 (brown cartoon/surface). The residues that were replaced by the insertion are shown as red sticks on the tralokinumab cartoon representation of the structure, while blue sticks show affinity-improving BAK1.1 substitutions (V_L_ CDR1 N27Y and V_H_ CDR3 N99S). (b) Both beneficial insertions are located on the periphery of the antigen binding site. The blue dotted line includes residues of the structural paratope of the BAK1.1 antibody. The colours indicate residues of the structural paratope, the positions of beneficial insertions and substitutions in yellow, red and blue, respectively. The variable heavy domains (V_H_) is highlighted in grey, while the variable light domain (V_L_) is shown in cyan.

The observation that loop extensions (possibly unstructured, see below) confer affinity enhancements is reminiscent of natural affinity maturation. CDR3 loop length variation can result from the V(D)J rearrangement leading to some length variation (and as such complicates the estimation of InDel frequency during somatic hypermutation of these loops). The identification of beneficial insertions in the V_L_ CDR3 of the BAK1 antibody in this study combined with the high tolerance in length variation of this loop in Nature suggests that length diversification of the V_L_ CDR3 might also represent a mechanism of affinity maturation. The 67aE insertion is located within the fourth loop in the third framework region (FWR3) (Fig. 6a), sometimes referred to as CDR4 (39). In contrast to the other hypervariable loops (CDR1-3), antibodies in Nature show minimal variation in this loop in the germline (51, 52). The antibody PGT121, which originated from the same germline gene (L-Vλ3-21*02) as the BAK1 V_L_, has a three amino acid insertion in the same position in the FWR3 as BAK1-INS1 (53). The frequent presence of insertions in the FWR3 loop in extensively affinity matured antibodies (anti-HIV-1 bnAbs) has been noted previously. For example, 5 of 17 antibodies with broad neutralization activity (29.4%) have a FWR3 loop insertion (29). These antibodies have been shown to confer broad neutralization activity of the 8ANC195 and 3BNC60 bnAbs (29, 30) and suggest that the framework region 3 is crucial for specificity determination.

The positions of both insertions in the FWR3 and V_L_ CDR3 loops on the tralokinumab:IL-13 complex suggested at least partial surface exposure and a position on the periphery of the antigen-binding site. The close proximity of the sites of both insertions to the antigen combined with their large effect on affinity suggests that it is possible that insertions induce direct interactions with the antigen.

The idea that insertions may exert their effects by making new contacts with the antigen is supported by studies of the role of insertions in affinity matured anti-HIV-1 bnAbs in Nature, which have shown that insertions extend loops and could form new intermolecular contacts, either directly or by enabling other loop residues flanking the position of the insertion. These could reach the antigen surface and interact with the antigen. Indeed, a FWR3 loop insertion of the anti-HIV-1 bnAb 8ANC195 (54) has been shown to exert its effect by extending the loop and forming intramolecular contacts. These in turn enable new contacts of FWR3 loop residues flanking the insertion with residues of the gp120 loops D and V5. The functional effect of the insertion was an extended epitope surface that included a pocket on the gp120 surface, resulting in broadened neutralization activity of the antibody against various viral strains. Another example is the four-residue insertion in the V_H_ CDR3 of the bnAb NIH45-46, which belongs to the VRC01 clonal lineage (55, 56). In bnAb NIH45-46 not only intra-molecular interactions but also new hydrogen bonds and electrostatic interactions of the insertion residues with the antigen are introduced by extending the CDR3 loop and, consequently, increasing the buried interface area. Apparently beneficial insertions can be found not only in the CDR loops, but also in a fourth loop in the FWR3.

The fact that BAK1 beneficial insertions are likely to have gained improved affinity by making new contacts with the antigen on the periphery of the interface, suggests that the improvements in affinity could be attributed to increased buried surface area of the antigen-binding site. Finally, in addition to static shape complementarity, insertions may change antibody affinity by changing the conformations of adjacent loops, as in the 2D1 anti-HA antibody insertion (57) at the junction of CDR2 and FWR3. This insertion altered the conformation of the CDR1 loop, resulting in 35-fold higher affinity. Overall, the mechanisms by which insertions exert their function include extension of loops to make new contacts with the antigen or alteration of the conformation of adjacent loops.

### Epistatic interactions and synergy of point substitutions in positions flanking the insertion site and InDels

Epistatic interactions between point substitutions are often governing the functional consequences of individual mutations, so that the combined effect of mutations is often not simply additive (7, 58). Even though epistasis was shown to be largely beneficial amongst CDR variants (59), mutations may still clash, requiring consensus design to avoid incompatible combinations e.g. in libraries generated by error-prone PCR (60). Whether the acquisition of beneficial *insertions* would depend on the presence of other point substitutions across the antibody variable region was not known before designing the insertion libraries. The discovery of insertions that improved BAK1 affinity in two different loops, independent of the presence of other point substitutions across the scFv gene, suggests that the effect of insertions does not necessarily depend on the presence of other mutations. This fact has important engineering implications, as it allows the identification of beneficial insertions without the need to sample large areas of sequence space with point substitutions, thus streamlining the search for beneficial insertions.

Although the effect of insertions did not depend on point substitutions far from the positions of the insertions, positive epistasis was observed with positions around the insertions. Insertion in the FWR3 (67aE) was beneficial only in the presence of a specific amino acid residue at position 68 (G68W), indicating epistatic interaction between those modifications, with a specific substitution at the position flanking the insertion and enabling the insertion to exert its function (i.e. permissive mutation). These results underline the importance of sampling high diversity of point substitutions: to identify an insertion variant with improved affinity, a combination of mutants flanking the position of the insertion with insertions of high diversity may be necessary to create a synergistic effect.

The idea that longer insertions generally lead to *higher* affinity, because an additional structural feature provides additional interactions is addressed in Fig. 5A. Apparently, longer insertions can, but do not have to lead to affinity maturation, so the specific context that determines strength of newly formed interactions matters.

Variants with insertions in the V_L_ FWR3 and the V_L_ CDR3 loops were found to have higher affinity for IL-13 but no ‘substitution-only’ variants in these loops were found to have conferred a beneficial effect on affinity. In contrast, variants with beneficial substitutions were found in different loops, namely the V_H_ CDR3 and V_L_ CDR1, in which no beneficial insertions were found. When these are combined, they act synergistically. This observation contrasts with the prior suggestion that most InDels share the *same* hotspots with point mutations because InDel duplications or deletions involve AGY motifs (21). In our selections, insertions and substitutions can have complementary mechanisms of function, meaning that insertions offer alternative routes to affinity maturation.

### A non-natural route to affinity maturation by InDel mutagenesis that mirrors natural length variation: TRIAD followed by insertional scanning mutagenesis (ISM)

Based on studies of the percentage of InDels in affinity matured antibodies in Nature, affinity-improving InDels were expected to be rare (21, 23, 24). While V(D)J rearrangements contribute to loop length variation (e.g. in the V_L_ CDR3), the contribution of InDels introduced by SHM *in vivo* is difficult to discern, although indels are evident as structural features. The mechanism by which insertions occur in Nature during SHM, by duplication of repeated sequences (21, 22), suggests that insertions with only limited diversity of amino acid replacements can be sampled by SHM. By contrast, our work directly introduces universal insertions and deletions across the entire gene and allows the construction of libraries of insertions that carry high amino acid diversity. We show the utility of this approach by successful affinity maturation caused by insertions in two different positions of the same antibody. Notwithstanding the question of whether and how indels are involved in SHM, the affinity improvements reported here result from changes in two different loops, the V_L_ CDR3 and the V_L_ FWR3. These positions in the CDR loops are selected for, as diversity was introduced across the entire variable domains of the BAK1 antibody: alternative positions would have been available, yet they were not selected. The importance of these positions for affinity is supported by the presence of minimal germline variation in the length of V_L_ FWR3 loop, at the position of the BAK1-INS1 insertion (51), suggesting that this loop can accommodate length variation. In a previous attempt to engineer antibody binding by increasing the length of the FWR3 loop (61), no affinity enhancement was achieved. Our work demonstrates that specific changes in the V_L_ FWR3 loop must be selected from *large* libraries and mirrors the high tolerance in length variation of this loop in Nature, making it the first successful combinatorial *in vitro* InDel engineering of an antibody.

### Strategy of InDel mutagenesis followed by targeted insertion (TRIAD-ISM)

This work presents the first evidence of a deliberate strategy in which insertions and point substitutions can have synergistic effects on affinity. The implication of this observation is that libraries used in selections would benefit from exploiting the effects of InDels and substitutions to explore additive effects on affinity. For protein engineering purposes the fact that both insertions and point substitutions can have complementary functions suggests that insertions can offer an alternative in cases where the identification of further beneficial point substitutions has proven difficult, thus increasing the chances of success in antibody evolution and offering additional pathways to affinity maturation. Combining multiple mutagenesis approaches leads to libraries that are too large to be screened comprehensively. Therefore, a strategy for combining mutagenesis approaches is necessary. Generally, point mutations can be introduced at any stage of this process, as long as the epistatic properties of the protein fold allow synergy of mutations. For the introduction of InDels we have explored in this work a two-step sequence that extends the simple introduction of InDels by TRIAD, where short (single) amino acid insertions are sampled entirely randomly. Given the experimental limits of screening, it is necessary to break down this procedure into two steps that answer the question where InDels are tolerated before the combinatorial diversity of the identity and length of potential insertions are addressed.

1. *Identification of InDel tolerance.* TRIAD is used to scan insertion points via single amino acid insertions by screening for binding or catalytic function. This outcome of this experiment will identify regions of the protein in which insertions are tolerated. Alternatively, insertion points can be deduced from previous knowledge of insertion tolerance (see discussion below).
2. *Insertional Mutagenesis.* In this semi-random approach, a library with insertion points based on the results of step (1) is made that contains high diversity insertions, i.e. with different lengths (in our case up to six amino acids) and amino acid diversity in each position of the inserts, followed by screening.

Step (1) can be achieved in an entirely combinatorial way: in contrast to previous methods of insertional mutagenesis (26, 62, 63), scanning allows prediction of positions that tolerate insertions not based on structural information, which is often not available, or natural length variation (which would indicate too many positions that can tolerate insertions, while scanning can indicate fewer positions that could be randomized). Previously high diversity indels have not been sampled at random positions, either because of restrictions in the diversity of indels that can be practically introduced (22) or because they only used rational instead of random approaches (26). Alternatively, prior knowledge can also be used as a guide instead of a random exploration by TRIAD. For example, the randomization of the V_L_ CDR3 of an anti-hapten antibody by insertional mutagenesis was based on the crystal structure of the complex to identify variants with modified specificity (63). Similarly, comprehensive mapping of fitness effects in response to InDels will suggest tolerant regions (64). Alternatively, comparative bioinformatic and phylogenetic analysis may provides insertion points for ISM. For database sequences with known function, the location and size of InDels may be indicative of functional switches (65). This approach has been used, albeit at much lower throughput, in the active site remodeling of related lactonases/phosphotriesterases (62).

More generally the identification and exploration of InDel diversity may give access to different trajectories *en route* to novel functional proteins that so far have not been explored. Insertions have different mechanism of action than substitutions: extending loops and making new contacts with the antigen are much more profound changes than typically introduced by single amino acid substitution. This idea is underlined by the observation of mutations in loops, where no beneficial substitutions had been found previously, suggesting different functional potential of InDels vs point substitutions. Indeed, loop extensions have resulted in introduction of additional specificities to create a multispecific ‘two-in-one antibody’ (66–70). Such versatile reagents offer the opportunity of neutralising multiple human targets for therapy (e.g. multiple, related cytokines). Similarly, making antibodies which hit both the human and rodent orthologues of a particular target would be desirable at the preclinical stage of drug development in order to understand the biology of the target in a different species. These approaches have been exemplified for point substitutions (66–70), but given the potential for introducing conformational changes *via* InDels, their potential in this respect may be explored in future work.

## Materials and Methods

### Construction of BAK1 libraries with random indels

BAK1 libraries with random indels throughout the entire scFv coding sequence were constructed using TRIAD as described previously (34, 71) *(SI Appendix*, Fig. S1 and S2). Briefly, the transposition reactions were performed starting from the vector pIDR9 (34) containing the BAK1 scFv coding sequence (pIDR9-BAK1) using either the TransDel or TransIns engineered transposon, to create libraries with deletions or insertions, respectively, and the enzyme MuA Transposase (Finnzymes). The cassettes Ins1, Del1, Del2 and Del3 were extracted from pUC57 plasmids (34). For each step, the ligation reactions were concentrated using the Zymo Research, DNA Clean & ConcentratorTM-5 and then transformed into * 30 μl of E. cloni^®^ 10G Electrocompetent Cells (Lucigen) (> 5 x10^9^ cfu/μg) by electroporation. The transformation efficiency was * 2 x 10^6^ colonies in each step. The libraries were subcloned into the ribosome display vector (33, 72) and the quality of the libraries was assessed by sequencing *(SI Appendix,* Table S1).

### Construction of the BAK1 InDel library Rec

DNA of the BAK1 libraries 3nt-Del, 6nt-Del, 9nt-Del and 3nt-Ins was mixed and recombined following a Staggered Extension Process *In-Vitro* DNA Recombination (SteP) protocol (38), using a high fidelity DNA polymerase (Phusion^®^ High-Fidelity DNA Polymerase, error rate 4.4 × 10^-7^) to keep the point mutagenesis as low as possible. The recombined BAK1 InDel library (Rec library) into the ribosome display vector was analysed by sequencing *(SI Appendix,* Table S2). This library (≈ 5×10^11^ variants, quantified by determining the concentration of plasmid DNA that was added into the transcription reaction, using the correlation 1 μg of 1200 bp DNA equals 7.5 x 10^11^ molecules) was the input of the first round of ribosome display selection. *In vitro* transcription, translation and selections were performed as described previously (33, 72), but using a high fidelity DNA polymerase for the amplification steps between the rounds of selection.

### Ribosome display selection from the Rec library

scFv-ribosome-mRNA complexes were subjected to selection in solution using recombinant biotinylated human IL-13 (Peprotech) to allow capture using streptavidin-coated paramagnetic beads (Dynabeads M-280), as described previously (35). Biotinylation of IL-13 surface lysine residues was done using EZ link NHS-LC-Biotin according to the manufacturer’s instructions (Pierce). Four rounds of K_D_-based selections were performed by panning with various biotinylated IL-13 concentrations below the K_D_ (< 2,300 pM). More specifically, the output of the selection with the lowest IL-13 concentration that gave a positive selection outcome (100 to 20 pM) was used in the following round of selection.

### Construction of libraries with insertions in the V_L_ CDR3

Insertional mutagenesis in the V_L_ CDR3 of the BAK1 scFv was achieved by generation of libraries in the phagemid vector pCANTAB6, as previously described (73), by oligonucleotide-directed mutagenesis. Five libraries were constructed using 5 different oligonucleotides designed to introduce randomized insertions of different lengths, from 0 to 5 amino-acid residues. Following the residue W91, blocks of 5 or 6 consecutive residues (NNS randomization) were randomized by Kunkel mutagenesis (74). Each library contained >10^9^ variants, estimated from the number of transformants after electroporation of *Escherichia coli* TG1 cells with the DNA product of each mutagenesis reaction.

### Phage Display selections of V_L_ CDR3 insertion libraries

Four rounds of K_D_-based selections of the BAK1 libraries with insertions in the VL CDR3 to enrich variants with improved affinity were performed in solution by capturing biotinylated human IL-13 with streptavidin-coated paramagnetic beads, following the principles described previously (75). Starting from an IL-13 concentration in the *K_D_* range of 10 nM in the first round, we decreased the antigen concentration by 10-fold in each round.

### Construction of BAK1 IgG variants by site-directed mutagenesis

Site-directed mutagenesis of BAK1 scFv coding sequence into pCANTAB6 was performed following the manufacturer’s instructions in the QuikChange Site-Directed Mutagenesis Kit (Agilent Technologies). The mutations introduced in the BAK1 or the BAK1-INS1 template sequences were: N99S in the V_H_ CDR3, N27I in V_L_ CDR1, G68W or G68E in the V_L_ FWR3. The BAK1 V_H_ and V_L_ sequences were then subcloned into the heavy chain (pEU1.3) and the λ light chain (pEU4.4) mammalian IgG1 expression vectors (76), respectively, which were designed as described previously (33).

### Expression and Purification of scFv and IgG variants

*E. coli* periplasmic extracts containing His6-tagged scFv variants of either the 3nt-Ins library or the outputs of phage display selections were produced for screening by HTRF assays (see *SI Appendix),* using the vector pCANTAB6 (73). scFv expression in *E.coli* was induced using isopropyl β-D-1-thiogalactopyranoside (IPTG). To induce cell lysis the cell pellets were resuspended in osmotic shock buffer MES (containing 50 mM MOPS, 0.5 mM EDTA and 0.5 M sorbitol; final pH 7.4).

His6-tagged scFv variants were purified for affinity-based screening using a competition HTRF assay. The expression of scFv variants in E. coli was induced using IPTG, while the cells were lysed using osmotic shock buffer TES (containing 200 mM Tris-HCl, 0.5 mM EDTA, 0.5 M sucrose; pH 8.0). The periplasmic fractions were purified on HisTrap™ HP (GE Healthcare) packed columns followed by buffer exchanged to PBS (NAP-10 columns; GE Healthcare).

IgGs were produced by co-transfection of the heavy chain and light chain pEU vectors (76) in mammalian cells and the IgG1 variants were purified by protein A affinity chromatography (MabSelect Sure columns, GE healthcare) using the AKTA express system and eluted with 0.1 M sodium citrate (pH 3.0). PD10 columns (GE healthcare) were used for buffer exchange of the antibodies in phosphate buffered saline (PBS) and the concentrations were measured by absorption spectrophotometry based on calculation of the extinction coefficient (ε) of each IgG variant based on their amino acid content.

## Supporting information

Skamaki_SI

## Acknowledgements

We thank Medimmune’s Biologics Expression Team (for assistance with protein production), their DNA Chemistry Group (for primer synthesis and sequencing) and Dr Mariana Rangel for preliminary crystallisation experiments. This research was funded by the Biological and Biotechnological Research Council (BBSRC) via grant BB/L002469/1 and ‘sparking impact’ funds, the EU Programme H2020 and MedImmune/AstraZeneca. KS received a BBSRC CASE studentship (BB/K012665/1) in collaboration with MedImmune/AstraZeneca. FH is an ERC Advanced Investigator (grant no. 695669).

## Abbreviations used

bnAbs: broadly neutralizing monoclonal antibodies
CDRs: complementarity determining regions
Fab: fragment antigen binding
FWR: framework regions
Indels: insertions and deletions
scFv: single-chain variable fragment
V_H_: Variable heavy chain domain
V_L_: Variable light chain domain

