## Supplementary material for "*In vitro* Evolution of Antibody Affinity via Insertional Mutagenesis Scanning of an Entire Antibody Variable Region": Skamaki_SI

### 1. BAK1 libraries with random indels for an affinity maturation campaign.

#### 1.1. TRIAD mutagenesis strategy.

BAK1 libraries with random indels throughout the entire scFv coding sequence were constructed using TRIAD (1) (Fig. S1 and S2). The BAK1 libraries 3nt-Del, 6nt-Del, 9nt-Del and 3nt-Ins contained 3, 6 and 9 nucleotide deletions and 3 nucleotide insertions (triplet NNN), respectively. The quality of the TRIAD libraries was assessed by sequencing (Table S1). The functional library sizes were at least 10-fold higher than the theoretical library sizes, suggesting that the percentages of variants with in-frame indels was sufficient to cover the theoretical diversity of each library. Very high (> 90%) percentage of variants with InDel at unique positions demonstrates that indels were randomly distributed across the entire scFv coding sequence and the gene is fully covered. In-frame indel variants with extra point mutations across the scFv sequence were not observed. As a consequence, epistatic interactions between indels and extra point substitutions should not affect the outcome of the selection, as deleterious or beneficial point substitutions would not mask the effect of beneficial indels. Overall, the quality control suggests that we successfully generated libraries with random InDels across the entire scFv gene, without extra point substitutions, and that this custom-made diversity in sequence space could be explored in library selections.

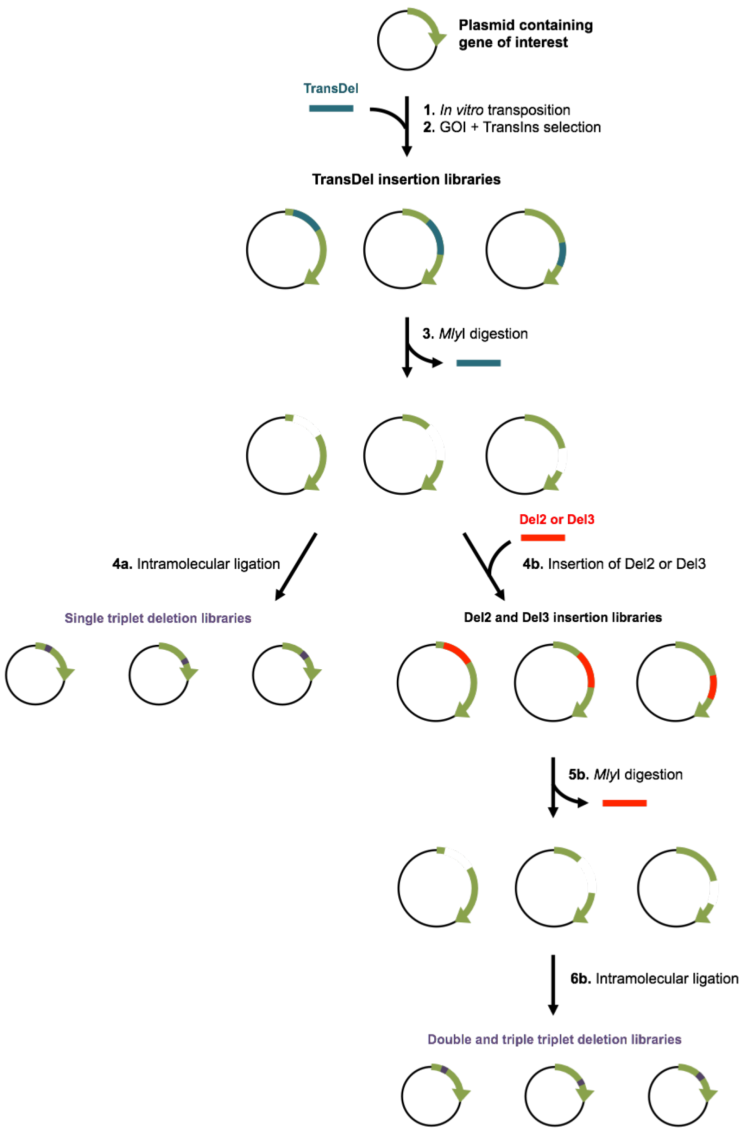

**Figure S1.** Schematic representation of the cloning steps followed for the construction of libraries with random 3-nt (2), 6-nt and 9nt deletions by TRIAD (1). Insertion of the transposon TransDel at random positions of the gene of interest is followed by removal of the transposon by digestion using the restriction site Mly at the flanking sequences of the engineered transposon. Intramolecular ligation of the library plasmids results in the deletion of a single triplet nucleotide, while insertion of a secondary cassette (Del2 or Del3) followed by MlyI digestion leads to removal of either double or triple triplet nucleotides. Copied with permission from (1).

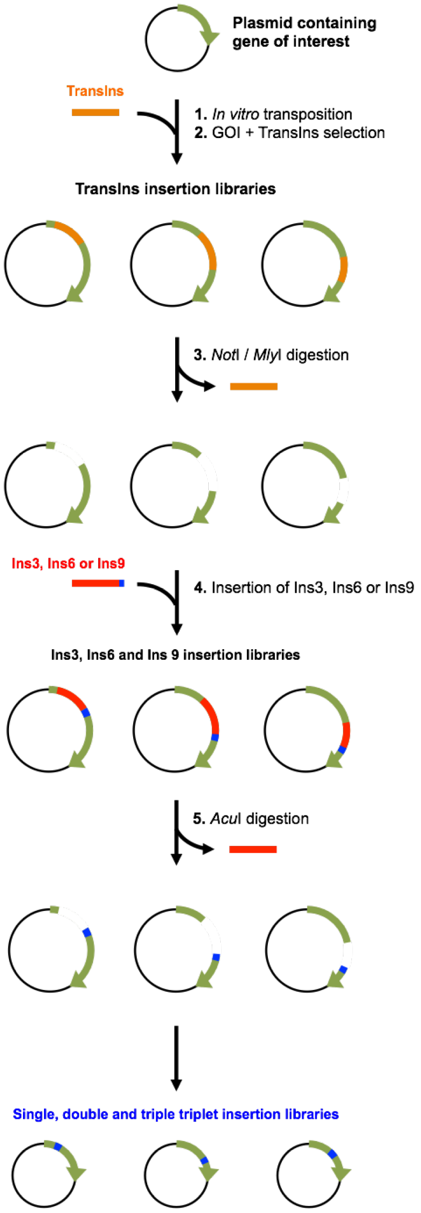

**Figure S2.** Schematic representation of the cloning steps followed for the construction of libraries with random 3-nt, 6-nt and 9nt insertions by TRIAD (Emond et al, 2018). Insertion of the transposon TransIns at random positions of the gene of interest is followed by removal of the transposon by digestion using the restriction sites MlyI/NotI at the flanking sequences of the engineered transposon. Insertion of a secondary cassettes (Ins3, Ins6 or Ins9), which contain NNN triplet insertions, followed by AcuI digestion leads to addition of single, double or triple triplet nucleotide insertions. Copied with permission from (1).

**Table S1.** Sequence analysis[a] of BAK1 libraries with random InDels.

| BAK1 InDel library | Variants with in-frame InDel of desired length % | Variants with frameshifts % | Parent (or linker) % | Variants with indel at unique position % | Theoretical library size | Functional library size |
|---|---|---|---|---|---|---|
| Number of nucleotide (nt) Insertion (Ins) / Deletion (Del) | Total | | | | | |
| 3 nt-Del | 86 | 14 | 0 | 90 | $7.5 \times 10^2$ | $1.7 \times 10^6$ |
| 6 nt-Del | 17 | 67 | 16 | 93 | $7.5 \times 10^2$ | $3.4 \times 10^5$ |
| 9 nt-Del | 29 | 52 | 19 | 83 | $7.5 \times 10^2$ | $5.8 \times 10^5$ |
| 3 nt-Ins | 36 | 47 | 17 | 100 | $4.8 \times 10^4$ | $7.1 \times 10^5$ |

[a] The analysis was based on sequencing of 88 variants from each library. The actual library size (≈ 2 x $10^6$ variants) was calculated by measuring the colony number after each cloning step. For each library, the functional library size shows the number of variants with in-frame InDel of desired length.

**A**

```
1   FWR1 10        20          30 CDR1   40  FW2    50 CDR2      60
EVQLVQSGAEVKKPGASVKVS-CKASGYT-FRNYGLSWVRQAPG-QGLEWMGWISAN-NG-DTNYGQE
                     V       LH             G            IY E

        70  FWR3   80            90      CDR3   100
FQG-RITMTT-ETSTNTAHME-LRSLRSDDTAVY-YCVRDSSSNWAR-WFF-DLWGKGTMVTVSS
   T      I          E            H            N   GY
```

**B**

```
1   FWR1  10       20   CDR1   30     FWR2  40        50 CDR2
SYV-LTQPPSVSVAPGQTARIPC-G-GNNIGSKLV-HWYQQ-K-PGQAPVLVVYDD-GDRPS
   L                   V T         G     Q RA           ES

   60 FWR3    70          80           90 CDR3         100
GIPERF-SGSNSGNT-ATL-TISRIDAGD-EA-D-YYCQVW-DT-GSDPV-VFG-GGTKLTVLG
      Y        DT  I         D  I Y      ES T     L   N
```

**Figure S3.** BAK1 library with random insertions across the scFv coding sequence. **(A)** Sequencing analysis showed that the insertions (red) were randomly distributed across the entire **(A)** $V_H$ and **(B)** $V_L$ regions (all variants have insertion at unique positions). Insertions or substitutions are shown in blue or red, respectively.

### 1.2. Rec library.

The library 'Rec' (Table S2) was generated by recombination of the BAK1 InDels libraries (3nt-Del, 6nt-Del, 9-nt Del and 3nt-Ins) using Staggered Extension Process In-Vitro DNA Recombination (SteP) (3). The functional library sizes of the variants with single ($1.4\times10^{11}$) or double ($4.6\times10^{10}$) in-frame InDels per gene were at least 10-fold higher than the theoretical library sizes ($5 \times 10^4$ and $1.26 \times 10^9$, respectively), suggesting that the Rec library covered the diversity not only of all possible BAK1 variants with single InDel, but also a high number of all possible BAK1 variants with double InDels per gene.

**Table S2.** Sequence analysis[a] of recombined BAK1 InDel library ('Rec library').

| | Percentage | Theoretical library size | Functional library size | Actual library size |
|---|---|---|---|---|
| **Parent** | 31 | | $1.6\times10^{11}$ | |
| **Single indel variants (in-frame)** | 27 | $5 \times 10^4$ | $1.4\times10^{11}$ | $5\times10^{11}$ |
| **Double indel variants (in-frame)** | 9 | $1.26 \times 10^9$ | $4.6\times10^{10}$ | |
| **Out-of-frame** | 33 | | | |

[a] The analysis was based on sequencing of 176 variants from the Rec library.The number of possible variants with single InDel was the sum of possible variants of each InDel library (see Table S1). The actual library size was determined by the concentration of plasmid DNA that was added in the in vitro transcription reaction (1 µg of 1200bp DNA is $7.5 \times 10^{11}$ molecules). The functional library size shows the number of in-frame variants (either parent or variants with single or double in-frame InDel per gene).

#### 1.3. Selection results.

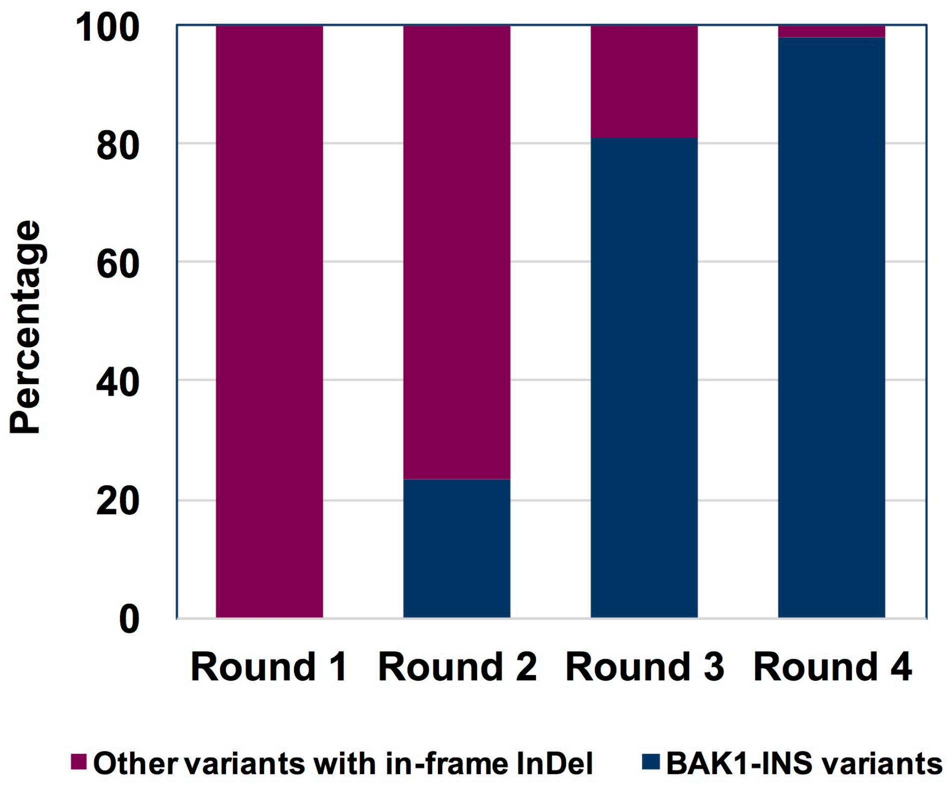

**Figure S4.** Ribosome display selections of the BAK1 InDel library Rec. A group of variants with different amino acid residue mutations at the same position, an insertion ($V_L$ 67a) and a point substitution ($V_L$ 68X), in VL FWR3 (BAK1-INS variants) were enriched, reaching 98% of the total variants with in-frame InDels in round 4.

The sequencing analysis of the ribosome selection outputs showed that, even though the percentage of BAK1 variants with point substitutions in round 4 was low, four variants with single point substitutions were enriched (Table S3). Two of these point substitutions had been found in a previous study (4) to confer large improvements on BAK1 affinity, BAK1.4 ($V_L$ CDR1 N27Y) and BAK1.55 ($V_H$ CDR3 N99S), the combination of point substitutions of these residues make up the variant BAK1.1, confirming that the results are in agreement with the past affinity maturation campaign.

**Table S3.** Variants with single point substitutions in round 4.

| **Loop** | **$V_H$ FWR2** | **$V_H$ FWR3** | **$V_H$ CDR3** | **$V_L$ CDR1** |
|---|---|---|---|---|
| **Residue** | **46** | **80** | **99** | **27** |
| **BAK1** | E | M | N | N |
| **BAK1.4** | E | M | N | Y |
| **BAK1.55** | E | M | S | N |
| **BAK1.56** | E | R | N | N |
| **BAK1.57** | K | M | N | N |

### 2. BAK1 libraries with insertions in the $V_L$ CDR3 for an affinity maturation campaign

#### 2.1. Selection results

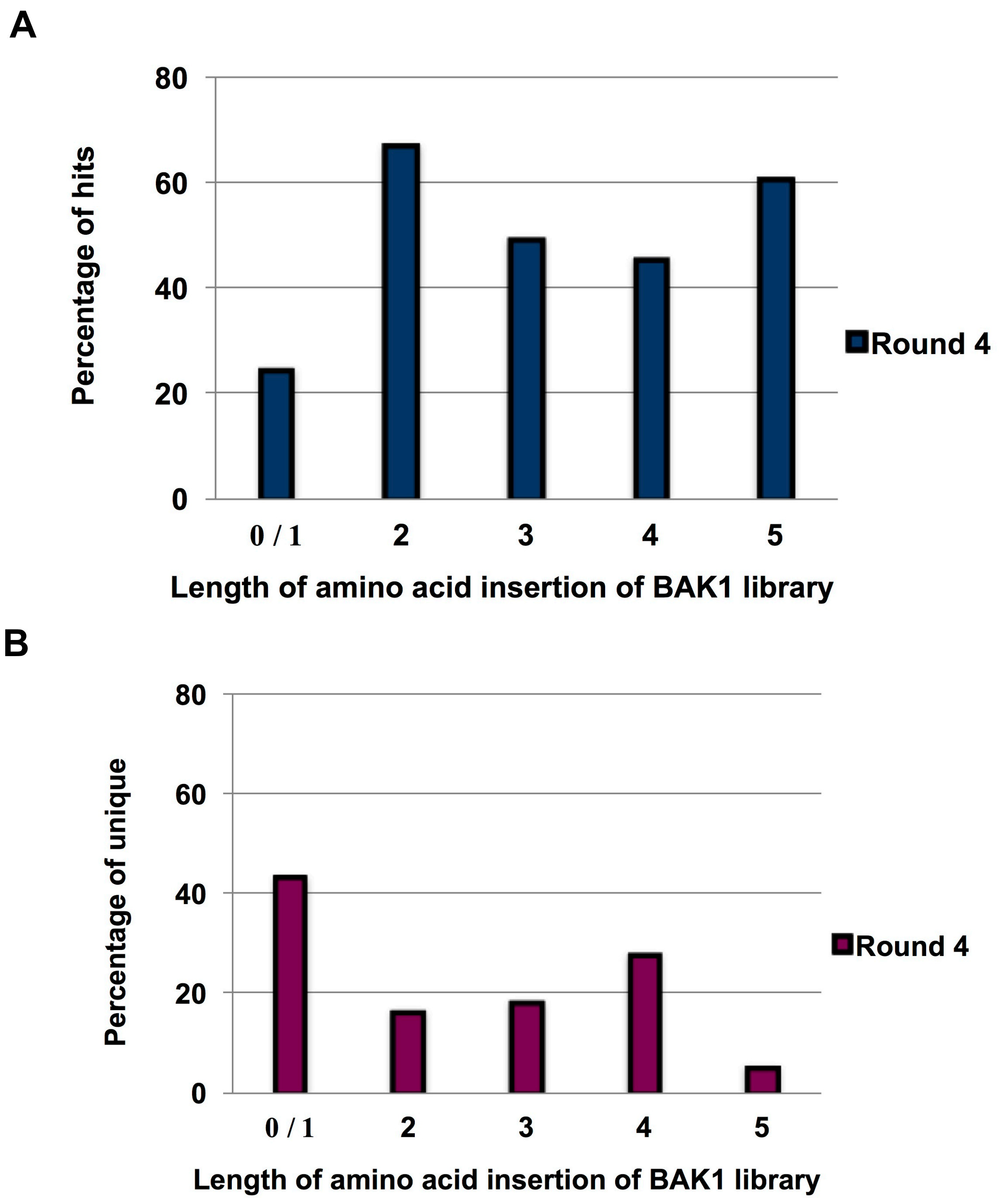

**Figure S5. Screening of the round 4 selection outputs for binding to IL-13.** 4 rounds of phage display selections of libraries with insertions in the $V_L$ CDR3 were followed by screening of 384 variants from the round 4 selection output of each library by an HTRF competition assay. Hits were considered the variants with signal same as the parent of higher in the HTRF assay. **(A)** The percentage of hits and **(B)** the percentage of unique hits were analysed by sequencing of all the hits for each library separately.

**Table S4.** Screening of BAK1 variants with insertions in the $V_L$ CDR3 for improved affinity.

| Loop | | VL CDR3 | | | | | | | | | | | Affinity | | |
|---|---|---|---|---|---|---|---|---|---|---|---|---|---|---|---|
| Residue | | 91 | 92 | 93 | 93a | 93b | 93c | 93d | 93e | 94 | 95 | 96 | $K_D$ (pM) Biacore | IC50 HTRF assay | IC50 ELISA (µg/ml) |
| Parent | BAK1 | W | D | T | - | - | - | - | - | G | S | D | 2,300 | 13.3 | 0.6 |
| | BAK1.1 | W | D | T | - | - | - | - | - | G | S | D | 50 | 0.3 | 0.06 |
| 1 | Ins1.2 | W | S | D | E | - | - | - | - | L | G | D | nd | - | 0.3 |
| | Ins1.3 | W | N | D | E | - | - | - | - | R | G | D | nd | 4 | 0.6 |
| | Ins1.4 | W | D | P | Y | - | - | - | - | R | G | D | nd | - | 0.6 |
| | Ins1.5 | W | A | G | D | - | - | - | - | K | G | D | nd | - | 3.9 |
| | Ins1.6 | W | S | W | H | - | - | - | - | G | A | D | 800 | 1 | 0.3 |
| | Ins1.7 | W | S | E | P | - | - | - | - | L | G | D | nd | 3.4 | nd |
| | Ins1.8 | W | N | W | A | - | - | - | - | D | G | D | nd | 8 | nd |
| 2 | Ins2.1 | W | S | G | W | D | - | - | - | P | G | D | 170 | 0.8 | 0.09 |
| | Ins2.2 | W | S | G | P | G | - | - | - | A | G | D | nd | 13.3 | nd |
| | Ins2.3 | W | P | G | P | M | - | - | - | R | G | D | nd | 4.2 | nd |
| | Ins2.4 | W | N | D | T | D | - | - | - | P | R | D | nd | 40 | nd |
| | Ins2.5 | W | R | T | W | D | - | - | - | S | G | D | nd | 0.7 | nd |
| | Ins2.6 | W | P | G | A | G | - | - | - | P | G | D | nd | 36.5 | nd |
| | Ins2.7 | W | A | D | G | Q | - | - | - | T | G | D | nd | 5.9 | nd |
| 3 | Ins3.1 | W | T | L | H | D | E | - | - | A | S | D | nd | 2.3 | nd |
| | Ins3.2 | W | T | A | D | F | P | - | - | P | S | D | nd | - | 5.2 |
| | Ins3.3 | W | P | G | P | D | P | - | - | V | S | D | nd | 20.9 | nd |
| | Ins3.4 | W | A | L | W | D | E | - | - | S | S | D | 300 | 0.4 | 0.03 |
| | Ins3.5 | W | P | N | W | E | V | - | - | G | S | D | nd | 0.9 | nd |
| | Ins3.6 | W | T | A | R | L | P | - | - | A | S | D | nd | 0.35 | nd |
| | Ins3.7 | W | V | P | W | G | S | - | - | H | S | D | nd | 1.2 | nd |
| | Ins3.8 | W | S | D | R | P | V | - | - | T | S | D | nd | 4.3 | nd |
| | Ins3.13 | W | A | G | P | L | Q | - | - | K | S | D | nd | nd | 1.7 |
| | Ins3.15 | W | P | G | P | D | A | - | - | L | S | D | nd | nd | 0.5 |
| | Ins3.17 | W | N | D | A | N | I | - | - | K | S | D | nd | nd | 0.9 |
| | Ins3.18 | W | N | P | F | D | T | - | - | R | S | D | - | - | 4 |
| 4 | Ins4.2 | W | V | P | P | H | I | R | - | G | S | D | 530 | 1 | 0.14 |
| | Ins4.3 | W | P | S | W | D | D | Q | - | G | S | D | 90 | 1.6 | 0.06 |
| | Ins4.4 | W | P | A | W | D | T | V | - | G | S | D | 34 | 1.3 | 0.07 |
| | Ins4.5 | W | V | P | G | P | Q | G | - | G | S | D | nd | 8.9 | nd |
| | Ins4.6 | W | V | G | P | E | H | L | - | G | S | D | nd | 5.2 | nd |
| | Ins4.7 | W | S | G | W | E | P | P | - | G | S | D | nd | 6.9 | 1.5 |
| | Ins4.9 | W | T | T | G | E | P | V | - | G | S | D | nd | nd | 1.5 |
| | Ins4.10 | W | T | T | F | D | Y | P | - | G | S | D | 360 | nd | 0.2 |
| | Ins4.11 | W | T | E | Q | T | A | K | - | G | S | D | nd | nd | 2.1 |
| | Ins4.12 | W | V | H | E | N | Q | N | - | G | S | D | - | nd | 0.1 |
| | Ins4.14 | W | V | S | K | V | E | E | - | G | S | D | nd | nd | 0.7 |
| | Ins4.15 | W | T | G | G | S | D | S | - | G | S | D | nd | nd | 0.9 |
| | Ins4.16 | W | V | D | P | S | N | P | - | G | S | D | nd | nd | 0.6 |
| | Ins4.18 | W | N | P | S | E | R | R | - | G | S | D | nd | nd | 2.7 |
| 5 | Ins5.1 | W | V | P | A | T | D | L | T | G | S | D | nd | - | 0.2 |
| | Ins5.3 | W | C | T | F | D | C | Q | T | G | S | D | 48 | 1.9 | 0.07 |
| | Ins5.4 | W | V | P | P | S | W | Q | T | G | S | D | nd | 0.9 | nd |
| | Ins5.5 | W | Y | D | R | A | E | I | T | G | S | D | nd | 3.8 | 0.3 |
| | Ins5.6 | W | V | A | E | D | P | P | T | G | S | D | nd | 2.7 | 0.2 |

### 2.2. Variants that tolerate insertions in the CDRs and the FWR3 loop

During the affinity maturation of BAK1 by insertional-scanning mutagenesis (ISM) 5,632 variants of the 3nt-Ins TRIAD library (in scFv format) with single amino acid insertions (Fig. 2A, Table S1) were screened with increasing stringency in an HTRF assay (see *3.2.1*). Initially 744 hits were identified based a low threshold (with a cut off of 10 for R1 and of 35 for R2), and then re-screened based on a more stringent cut-off (of 49 for R1 and of 70 for R2). Variants whose sequence analysis showed insertions in unique positions were screened in four repeats (to ensure genuine positives) and the Table below (and Fig. 4 in the main text) show the locations of insertions.

**Table S5.** Hits from affinity maturation of BAK1 by insertional-scanning mutagenesis (ISM).

| Region | | Insertions tolerated |
|---|---|---|
| $V_H$ | FW1 | -[a] |
| | FW2 | -[a] |
| | FW3 | I76a |
| | FW4 | -[a] |
| | CDR1 | - |
| | CDR2 | 5 x G53a ,<br>K55aS56 |
| | CDR3 | - |
| $V_L$ | FW1 | -[a] |
| | FW2 | -[a] |
| | FW3 | Y66aT67<br>2 x T67a<br>L67a<br>E67a |
| | FW4 | -[a] |
| | CDR1 | - |
| | CDR2 | - |
| | CDR3 | 4 x V91a<br>Q92aT93, P92aT93, S92aG93, N92aS93,<br>I92aA93, N92aS93, S92aT93, S92aT93, N92aT93, A92aT93<br>V93a, 2 x T93a, D93a, 2 x L93a, I93a, F93a, 2 x V93a, T93a,<br>S93a, L93a, I93a, I93a, I93a, I93a, D93a<br>Q96a97S, Q96a<br>A97a |

[a] Some insertions were found in FWR 1-4 in the experiments displayed in Figure 4.

### 2.3 Measurement of temperature stabilities of selected mutants

Thermal stability measurements of the improved IgG variants were carried out by differential scanning fluorimetry (DSF) (Fig. S6) (5). IgGs have two unfolding transitions, the unfolding of the CH2 domain is followed by that of the antigen-binding fragment (Fab) and the CH3 domain (6, 7). DSF does not resolve these two transitions, but for the IgG1 variants with the lowest $T_m$ values two peaks were observed: the first peak showing how the $T_m$ of the IgG is affected by changes in the variable regions of the Fab.

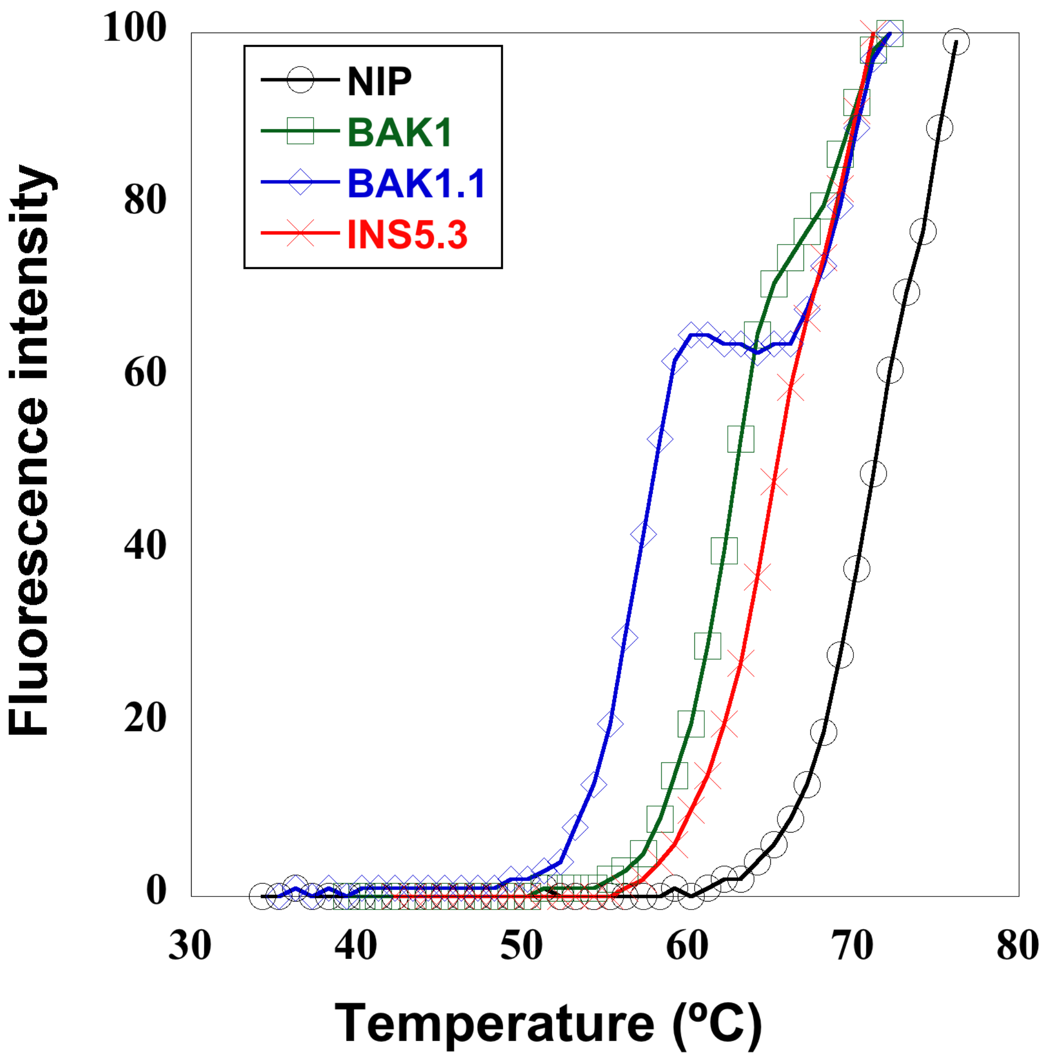

**Figure S6.** Thermal stability measurements of improved BAK1 IgG variants with insertions in the $V_L$ CDR3. The normalized melting curves are shown as measured by differential scanning fluorimetry (5). Measurements were repeated twice for all mutants.

### 3. Supplementary Methods

#### 3.1 Thermal stability measurements.

To determine the melting temperatures ($T_m$) of the BAK1 variants with improved affinity we used Differential Scanning Fluorimetry (DSF), using SYPRO Orange fluorescence to monitor unfolding of the IgG1 variants (5). 20 µL of IgGs with concentrations 0.75- 1.7 mg/ml were mixed with 5 µL SYPRO Orange dye (Invitrogen) at a final concentration of 625x. The fluorescence intensity was recorded over a temperature range of 25 °C to 99 °C in duplicate using a real-time PCR machine. The software GraphPad Prism software (GraphPad, La Jolla, CA) was used to determine the melting temperatures ($T_m$) by fitting the data to the following Boltzmann equation:

$$Y = Bottom + (Top - Bottom) / (1 + exp (T_m – X /Slope)$$

Here, Y represents the fluorescence intensity, X is the temperature; bottom and top are the minimum and maximum intensities, slope defines the slope of the curve within $T_m$ and $T_m$ is the melting temperature of the IgG variant.

**3.2. Binding assays and affinity measurements.**

**3.2.1 Homogenous Time Resolved Fluorescence (HTRF) Assays**

An HTRF binding assay was used to identify the variants of the 3nt-Ins BAK1 library that retained binding to IL-13. Screening was carried out using *E. coli* periplasmic extracts containing $His_6$-tagged scFv variants. 5,632 variants were tested in Corning® 384-well low volume non-binding plates. The assay buffer was composed of 0.1% bovine serum albumin, 0.4 M potassium fluoride and 1 x PBS. Each well contained 20 µL of the following reagents diluted in assay buffer: 1 nM streptavidin cryptate (CisBio), 5 nM biotinylated IL-13, 20 nM anti-his-XL665 (CisBio) and a 9-fold dilution of periplasmic fraction ($His_6$-tagged scFv).

An HTRF *competition* assay was used for screening of 384 variants from round 4 phage selection outputs of each library with insertion in the BAK1 $V_L$ CDR3. Each well contained 20 µL of the following reagents diluted in assay buffer to the following final concentrations: 1 nM Eu-labelled streptavidin cryptate (CisBio), 1.25 nM biotinylated IL-13, 1 nM Dylight 650-labelled BAK1-IgG1 and a 9-fold dilution of periplasmic fraction ($his_6$-tagged scFv). For further affinity-based screening the same assay was used to determine the IC50s of purified $His_6$-tagged scFv variants by titration of the scFv with two-fold serial dilutions, starting from 100 nM as the highest concentration.

For both assays the reaction mixture was incubated at room temperature for 4 hours to reach equilibrium and then read on a Perkin Elmer Envision plate reader, using a 320 nm excitation filter and 590 and 665 nm emission filters. The HTRF ratio was calculated by Ratio = (665 nm emission/620 nm emission) x 10,000. Non-specific binding was determined by calculation of the HTRF ratio in control wells. Specific binding was calculated according to

$$\Delta F = (\text{sample ratio} - \text{negative control ratio/ negative control ratio}) \times 100$$

#### 3.2.2 Competition ELISA

A competition ELISA was carried out to identify BAK1 IgG1 variants with enhanced affinity from the libraries with insertions in the $V_L$ CDR3. 96-well Nunc MaxiSorb flat-bottom plates were coated with BAK1-IgG1 in PBS (10 µg/ml) and incubated overnight at 4 ºC. The wells were blocked with PBS/3% milk and washed with PBS after 1 hour at room temperature. 50 µL of the mix of a two-fold serial dilution of the BAK1-IgG (30 µL, starting from 10 µg/ml, final 5 µg/ml) with IL-13-bio (30 µL, 0.2 µg/ml, final 0.1 µg/ml) in PBS-3% milk was incubated at room temperature while shaking for 1 hour. After washing with PBS/0.01% Tween binding was detected using a streptavidin conjugated to HRP antibody. Colorimetric detection was done by the addition of 50 µl of the substrate 3,3′,5,5′-tetramethylbenzidine (Sigma) and the reaction was stopped with 50 µl 0.5 M $H_2SO_4$.

#### *3.2.3 Surface Plasmon Resonance.*

$K_D$ measurements of the IgG1 variants were performed using the Biacore T200. Biosensor affinity measurements were performed using protein G-mediated capture immobilisation of the human antibodies and flowing the recombinant human IL-13 (PeproTech) as the analyte, as previously described (Groves et al., 2014), in the buffer HBS-EP+ (containing 0.01 M HEPES, 0.15 M NaCl, 3 mM EDTA, 0.05% v/v Tween 20; pH 7.4). A sensor surface pre-immobilized with a recombinant Protein G variant, the Series S Sensor Chip Protein G (GE Healthcare), was used for binding of the human IgG1 variants. IgG1 analysis was performed first by capturing ~100 resonance units (RUs) of the antibody on the protein G sensor chip (60 s capture at 5 µl/min) and running a two-fold serial dilution of recombinant IL-13, starting from a concentration of 6.25 nM with an association time of 5 min at 50 µL/min flow rate. Depending on the strength of the interaction, dissociation was followed for 300 -1800 sec. Regeneration of protein G surfaces was achieved by two consecutive 20 sec injections of 6 M guanidinium hydrochloride in PBS. The errors of the $k_{on}$ and $k_{off}$ measurements were between 0.01- and $0.09 \times 10^6\ M^{-1}s^{-1}$ and between 0.001 and $0.34 \times 10^{-4}\ s^{-1}$, respectively.
